# Pten, Pi3K and PtdIns(3,4,5)P_3_ dynamics modulate pulsatile actin branching in *Drosophila* retina morphogenesis

**DOI:** 10.1101/2023.03.17.533017

**Authors:** Jacob Malin, Christian Rosa Birriel, Victor Hatini

## Abstract

Epithelial remodeling of the *Drosophila* retina depends on the pulsatile contraction and expansion of apical contacts between the cells that form its hexagonal lattice. Phosphoinositide PI(3,4,5)P_3_ (PIP_3_) accumulates around tricellular adherens junctions (tAJs) during contact expansion and dissipates during contraction, but with unknown function. Here we found that manipulations of Pten or Pi3K that either decreased or increased PIP_3_ resulted in shortened contacts and a disordered lattice, indicating a requirement for PIP_3_ dynamics and turnover. These phenotypes are caused by a loss of protrusive branched actin, resulting from impaired activity of the Rac1 Rho GTPase and the WAVE regulatory complex (WRC). We additionally found that during contact expansion, Pi3K moves into tAJs to promote the cyclical increase of PIP_3_ in a spatially and temporally precise manner. Thus, dynamic regulation of PIP_3_ by Pten and Pi3K controls the protrusive phase of junctional remodeling, which is essential for planar epithelial morphogenesis.

**Graphical Abstract: Control of contact length by Pi3K, Pten and PIP_3_.:** 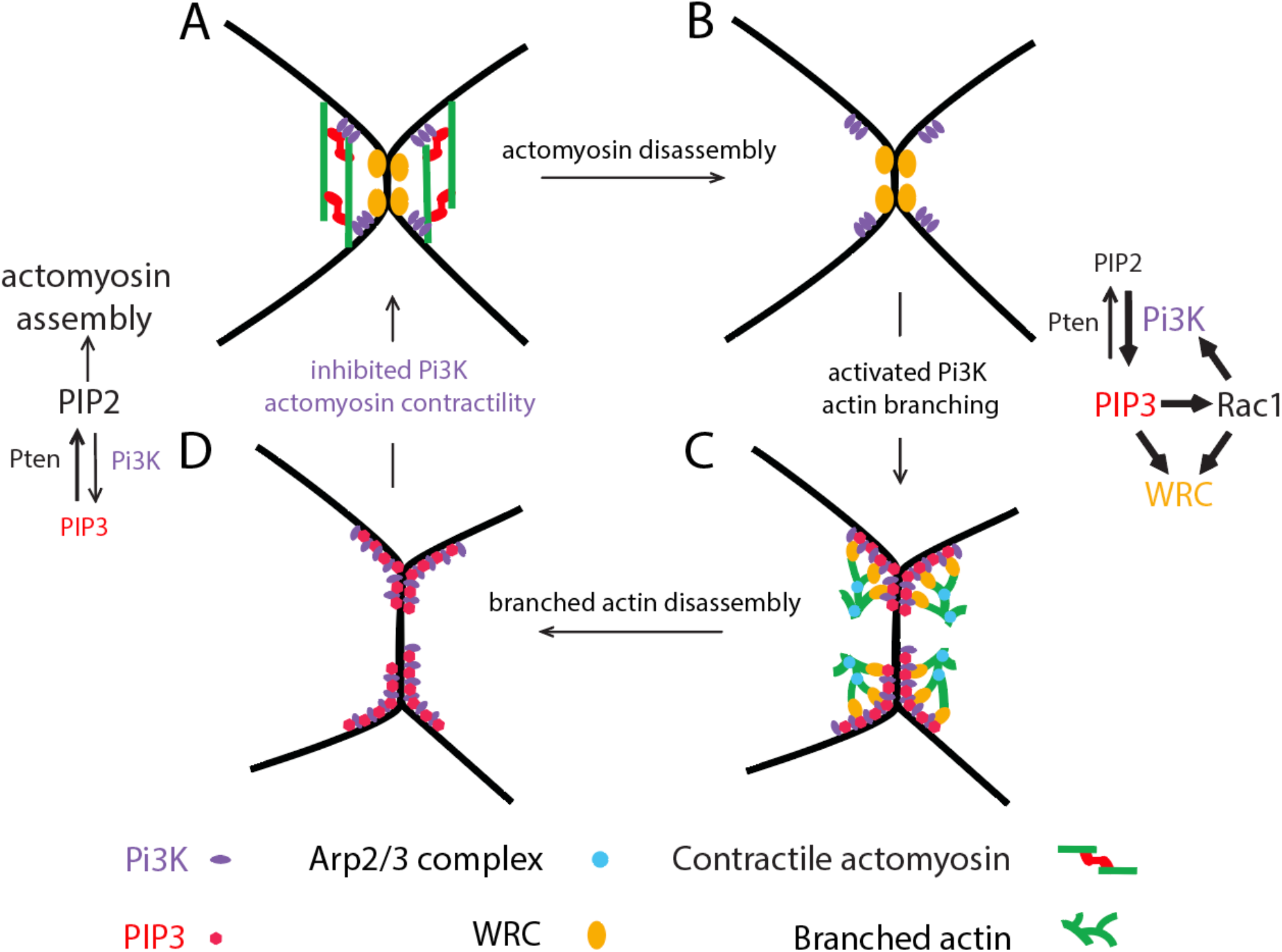

Pi3K regulates the transition from contraction to expansion through its tension-dependent localization to tAJs and modulation of its lipid phosphatase activity. Pten localizes uniformly to regulate PIP_3_ turnover and attenuate PIP_3_ production. (A) Tension shortens contacts, concentrates Pi3K at four spots at a distance from tAJs, and inhibits Pi3K’s lipid phosphatase function. (B) High tension ultimately disassembles contractile networks allowing Pi3K to flow toward tAJs, produce PIP_3_ and activate the WRC to promote actin branching and contact expansion. (C) High protrusion in expanded contacts disperses the WRC and disassembles the branched actin network. (D) Branched actin disassembly allows the assembly and contraction of an actomyosin network, which increases tension and contracts the contact leading to the flow of Pi3K away from tAJs and inhibition of its lipid phosphatase function, thus completing the cycle.

## INTRODUCTION

The *Drosophila* retina is a model for understanding epithelial morphogenesis (Cagan and Ready, 1989). The retina’s apical epithelium is composed of approximately 800 ommatidia containing stereotypically shaped and arranged cell types (Cagan, 2009; Carthew, 2007; Johnson, 2021). Pulsatile changes in cell junction length, depending on both contractile actomyosin and protrusive branched actin networks, are essential to ommatidial morphogenesis (Blackie et al., 2021; Blackie et al., 2020; Chan et al., 2017; Del Signore et al., 2018; Galy et al., 2011; Hayashi and Carthew, 2004; Letizia et al., 2019; Malin et al., 2022; Zallen et al., 2002). The role of contractile actomyosin pulsing has received considerable attention in similar systems (Heer and Martin, 2017). More recently understood is the role of protrusive forces as cell contacts expand, regulated by the WAVE regulatory complex (WRC), which activates the Arp2/3 complex to promote actin branching (Del Signore et al., 2018; Rottner et al., 2021). Inhibition of either actin branching or actomyosin contractility impairs this balance between expansion and contraction, leading to changes in cell shape and arrangement (Del Signore et al., 2018). Thus, a major goal is to determine the mechanisms that cyclically activate and inactivate cytoskeletal networks that govern both protrusion and contraction.

During eye epithelial remodeling, cell contacts cycle through contractile and protrusive pulses. A central player in this process is the homophilic adhesion molecule Sidekick (Sdk) (Letizia et al., 2019; Malin et al., 2022). Sdk dynamically associates with tAJs, where it toggles between interacting with contractile and protrusive effectors. To expand junctions, Sdk binds the WRC, which promotes actin branching and protrusion. As the contacts expand, Sdk exchanges interactions with the WRC for Polychaetoid (Pyd), the fly Zonula Occludens-1 (ZO-1) protein. Pyd then connects the junctional actomyosin networks to the tAJs to contract the junctions. As tension rises and contacts contract, Sdk exchanges interactions with Pyd for the WRC to initiate the next expansion cycle.

While we know that Sdk recruits the WRC, it is likely that additional coincident signals activate the WRC to generate protrusion. We previously discovered that the phosphoinositide PtdIns(3,4,5)P_3_ (PIP_3_) accumulates at tAJs of expanding cell contacts (Del Signore et al., 2018). *In vitro,* PIP_3_ synergizes with other signals to stimulate WRC-dependent actin branching, suggesting that it could regulate contact expansion (Chen et al., 2017; Hume et al.; Koronakis et al., 2011; Mendoza, 2013; Mendoza et al., 2011; Schaks et al., 2018). In this context, Rac1 controls the allosteric activation of the WRC, and PIP_3_ can activate Rac1 through intermediary Rac GTPase exchange factors (Cantley, 2002).

PIP_3_ levels are regulated by the phosphatase and tensin homolog (Pten) and the class 1A phosphoinositide 3-kinase (Pi3K), which respectively decrease and increase its levels through dephosphorylation and phosphorylation, converting it to and from PtdIns(4,5)P_2_ (PIP_2_) (Auger et al., 1989; Li et al., 1997; Traynor-Kaplan et al., 1988; Whitman et al., 1988). Pten and Pi3K are highly studied proteins that regulate cell growth, metabolism, and survival, and influence cancer progression (Gao et al., 2000; Goberdhan et al., 1999; Huang et al., 1999; Oldham et al., 2002). In these processes, they have multiple known effectors, with Pten dephosphorylating multiple proteins in addition to PIP_3_ (Chalhoub and Baker, 2009; Qi et al., 2020). Thus, it is important to analyze the relationship between these actors and their mechanism of action. While epithelial cell contacts are shortened in *pten* mutants, the importance of Pten to protrusive forces has never been examined (Bardet et al., 2013).

In the regulation of F-actin dynamics, Pten, Pi3K and PIP_3_ often exhibit highly specific subcellular distribution. For example, high Pi3K and PIP_3_ levels promote protrusion at lamellipodia of motile cells, while high Pten and PIP_2_ levels promote contraction at the rear and sides (Funamoto et al., 2002). Likewise, in certain epithelia, Pten associates with a protein located at the subapical AJs (Bazooka/Par3) and produces PIP_2_, which specifies an apical identity while PIP_3_ specifies basolateral identity (von Stein et al., 2005). Selective recruitment of Pi3K to edges of newly formed AJs has also been shown in vertebrates (Kovacs et al., 2002; Perez et al., 2008; Yamada and Nelson, 2007). However, in junction remodeling during planar epithelial morphogenesis, the relative localization of Pten, Pi3K and PIP_3_ has never been fully elucidated and the interactions between these factors are therefore unclear.

This study addresses this open question by examining both protrusive and contractile cytoskeletal elements and probing the function of Pten, Pi3K, and Rac1 in relation to PIP_3_. We show that Pten and Pi3K dynamically modulate PIP_3_ as well as cytoskeletal and mechanical forces that shape the epithelium, and through live imaging, reveal the subcellular organization of this system over time. We also provide evidence that PIP_3_ dynamics modulate Rac1 function, that Rac1 amplifies Pi3K function, and that both PIP_3_ and Rac1 activate the WRC, actin branching, and contact expansion. We propose that PIP_3_ dynamics regulate contact length pulsing by controlling the cytoskeletal transition from contraction to expansion.

## RESULTS

### *pten* controls epithelial remodeling of the retina

To examine *pten’s* role in eye epithelial remodeling, we generated eyes that were entirely mutant for three independent *pten* mutant alleles: *pten^3^*, *pten^c494^* and *pten^2L100^* (Devergne et al., 2014; Goberdhan et al., 1999; Oldham et al., 2002; Stowers and Schwarz, 1999). Consistent with previous reports, our data below indicate that *pten^3^* and *pten^c494^* are strong alleles while *pten^2L100^* is a hypomorph (Goberdhan et al., 1999; Huang et al., 1999; Oldham et al., 2002). In wild-type (WT), ommatidia form a regular hexagonal array with a small variation in edge length of each hexagon (Fig. 1A-B’, D). Scanning electron microscopy revealed that, in contrast to the regular shape and tessellation of ommatidia in WT adults, a fraction of ommatidia in *pten* mutant eyes were irregularly shaped and some were fused to one another (Fig. 1C-C’). To determine the effects of *pten* loss on epithelial organization, we examined eyes at 36-42h after puparium formation (APF) after cells acquired stable shapes (Fig. 1E-G). In *pten^3^, pten^c494^* and *pten^2L100^* mutants, ommatidia maintained their structure but formed quadrilaterals and pentagons in addition to hexagons (Fig. S1A). Defects in cell arrangements included clustering of multiple mechanosensory bristles (Fig. 1F orange arrowheads, Fig. S1B) and clustering of lattice cells (LCs) around bristles, resulting in cellular “rosettes”, a phenotype that can arise from impaired intercalation (Fig. 1E-G yellow arrowheads, Fig. S1C) (Bao and Cagan, 2005; Johnson et al., 2011; Johnson et al., 2008; Letizia et al., 2019; Malin et al., 2022; Seppa et al., 2008). In addition, some 3° LCs failed to occupy the corners of ommatidia, while others were missing, with remaining LCs occupying both the lattice edge and corner (Fig. 1E-G blue arrowheads, Fig. S1D). Also common was the partial or complete loss of LC-LC contacts and formation of contacts between 1° cells of adjacent ommatidia (Fig. 1E-G magenta arrowheads, Fig. S1D).

**Figure 1:**
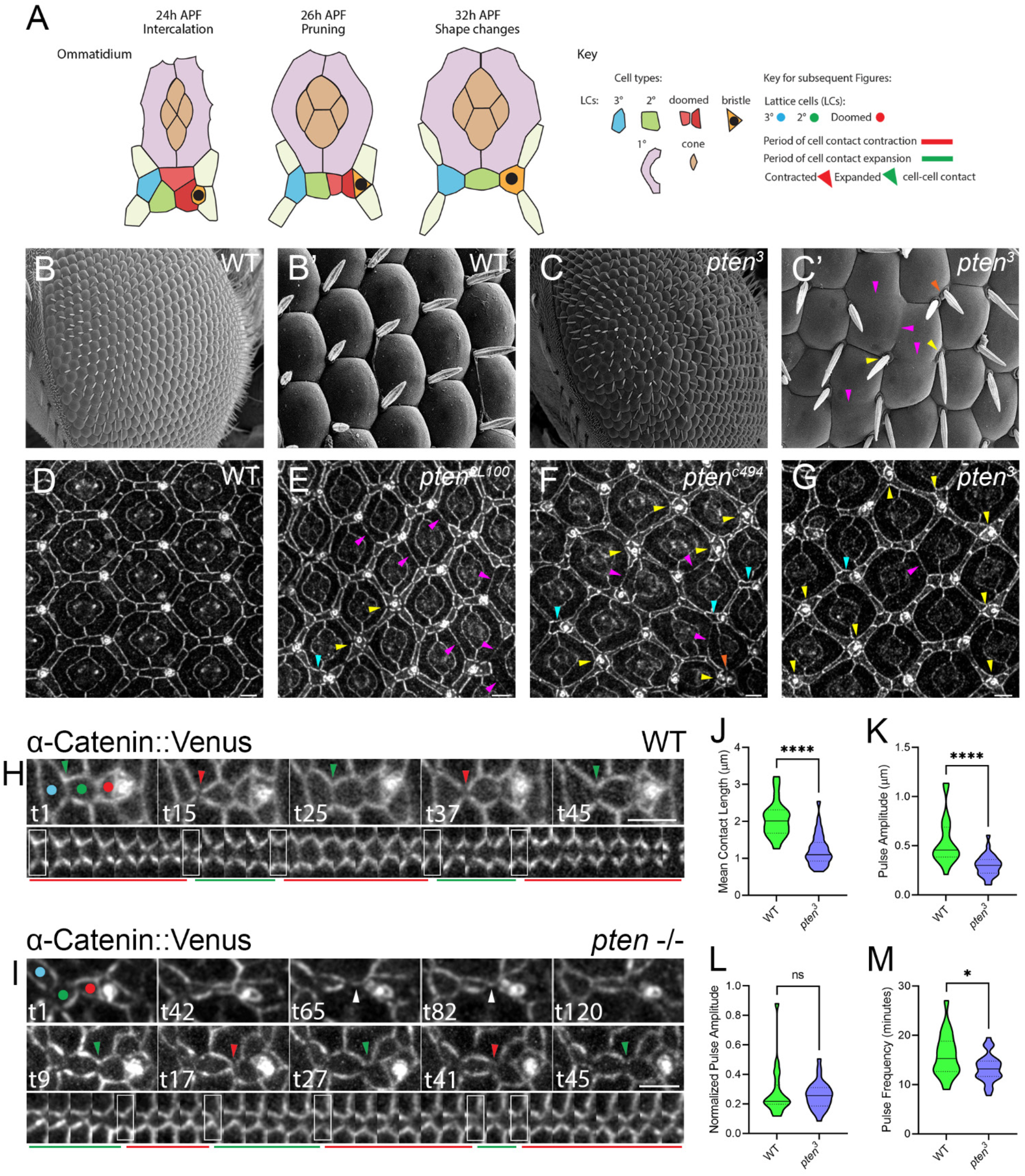
*pten* affects contact pulsing, epithelial remodeling and ommatidial structure. (A) Left: Major steps in lattice remodeling. Secondary (2°), tertiary (3) and mechanosensory bristle cells surround cone and primary (1°) cells. Right: Types of LCs, periods of expansion and contraction and contracted or expanded LC-LC contacts in this and subsequent figures. (B-C) Ommatidia in adult (B-B’) WT or (C-C’) *pten* mutant eyes. (D-G) Cellular defects in different *pten* mutant alleles compared with WT, 36-40h APF. (D) WT; each bristle and 3° LC connects to three 2° LCs. (E) *pten^2L100^*; common defects include cellular rosettes (yellow arrowheads) and LC-LC contacts lost and replaced by 1°-1° contacts (magenta arrowheads). (F-G) *pten^3^* and *pten^c494^*; defects are similar to *pten^2L100^* but more severe. Refer to Fig. S1A-D for quantification of specific defects. (H) Snapshots of a lattice edge and a kymograph of a single LC-LC contact at 2-minute intervals in a WT eye, 26-28 AFP. When a cell prunes from the lattice, a new contact is immediately established between adjacent cells. LC-LC contacts expand and contract repeatedly over periods of 10-20 minutes. (I) Cellular dynamics in a *pten* eye. Above: Snapshots of a LC being pruned from the lattice. After its delamination, adjacent cells remain separated, creating a contact between two 1° cells (white arrowheads) that is repaired over the next hour. Below: Snapshots and kymograph of a pulsing LC-LC contact. Compared with WT, expansions and contractions are small and uncoordinated, and the contact remains short. (J-M) Quantifications of contact length pulses in WT and *pten* mutant eyes. (J) Mean contact length. (K) Pulse amplitude. (L) Pulse amplitude normalized to mean contact length. (M) Pulse frequency. Statistics were performed using a Mann-Whitney *u* test, n=20 WT, 30 *pten* contacts from 2-3 eyes. *p>.05, ****p>.0001. Scale bars 5 μm in this and subsequent figures.

To determine whether *pten* acts autonomously, we generated marked *pten^3^* and *pten^2L100^* mutant cells and asked whether the cellular defects were confined to the clones (Fig. S1E-G). We found that the characteristic intercalation and cellular arrangement defects were largely confined to the *pten* mutant clones indicating that *pten* acts autonomously. We also found defects adjacent to the clones at a lower frequency. These autonomous and non-autonomous defects suggest that *pten* controls force dynamics and transmission necessary for epithelial remodeling. As the remodeling defects in *pten^3^* and *pten^c494^*eyes were more severe compared with *pten^2L100^* eyes, we used the *pten^3^* allele in subsequent experiments.

### *pten* controls fluctuations of LC-LC contact length

During lattice remodeling, the LC-LC contacts expand and contract repeatedly. These dynamics arise from the alternate generation of protrusive and contractile forces at the level of individual LC-LC contacts (Del Signore et al., 2018). When LCs are pruned, their neighbors rapidly reestablish contact (Fig. 1H, Movie 1). To determine whether *pten* affects these dynamics, we live imaged *pten* mutant eyes expressing α-Catenin::Venus to observe cell outlines at the level of AJs. We observed minimal expansion and contraction of LC-LC contacts and temporary or permanent 1°-1° contacts formed upon cell pruning and sometimes spontaneously (Fig. 1I, Movie 1). We then measured the length of LC-LC contacts over time. We excluded contacts that failed to expand following contraction and those that were being lost and replaced with 1°-1° contacts. We found that in *pten* mutants, the mean length, pulse amplitude, and pulse frequency of the LC-LC contacts decreased significantly compared with WT (Fig. 1J-K, M). However, the pulse amplitude normalized to average contact length was not significantly different from WT, indicating that the shortened contacts in *pten* mutants were still pulsing (Fig. 1L). These contact pulsing defects suggest that *pten* plays a role in rebalancing the opposing contractile and protrusive forces that contract and expand the LC-LC contacts.

### *pten* controls protrusive and contractile dynamics of LC-LC contacts

To better understand how Pten controls cytoskeletal dynamics, we compared F-actin and MyoII dynamics between WT eyes and *pten* mutant eyes. In time-lapse movies, we measured the length of LC-LC contacts and signal intensities of F-actin at the vertices and MyoII along the entire contacts where they concentrate most strongly. We detected F-actin using the actin-binding domain of Utrophin tagged with GFP (Utr::GFP) and MyoII with Spaghetti Squash (Sqh), the regulatory light chain of non-muscle Myosin II, tagged with GFP (Sqh::GFP) (Fig. 2A-D, Movies 2). We then computed the time-resolved Pearson’s cross-correlation between the signal intensity of either F-actin or MyoII and contact length (Fig. 2E-H). In WT, protrusive F-actin accumulated during contact expansion and peaked just prior to maximal expansion, while MyoII accumulated during contact contraction and peaked just prior to maximal contraction, consistent with previous reports (Del Signore et al., 2018; Malin et al., 2022). In *pten* mutant eyes, the positive correlation between F-actin accumulation and contact length decreased significantly compared with WT (Fig. 2E-F, I).

**Figure 2:**
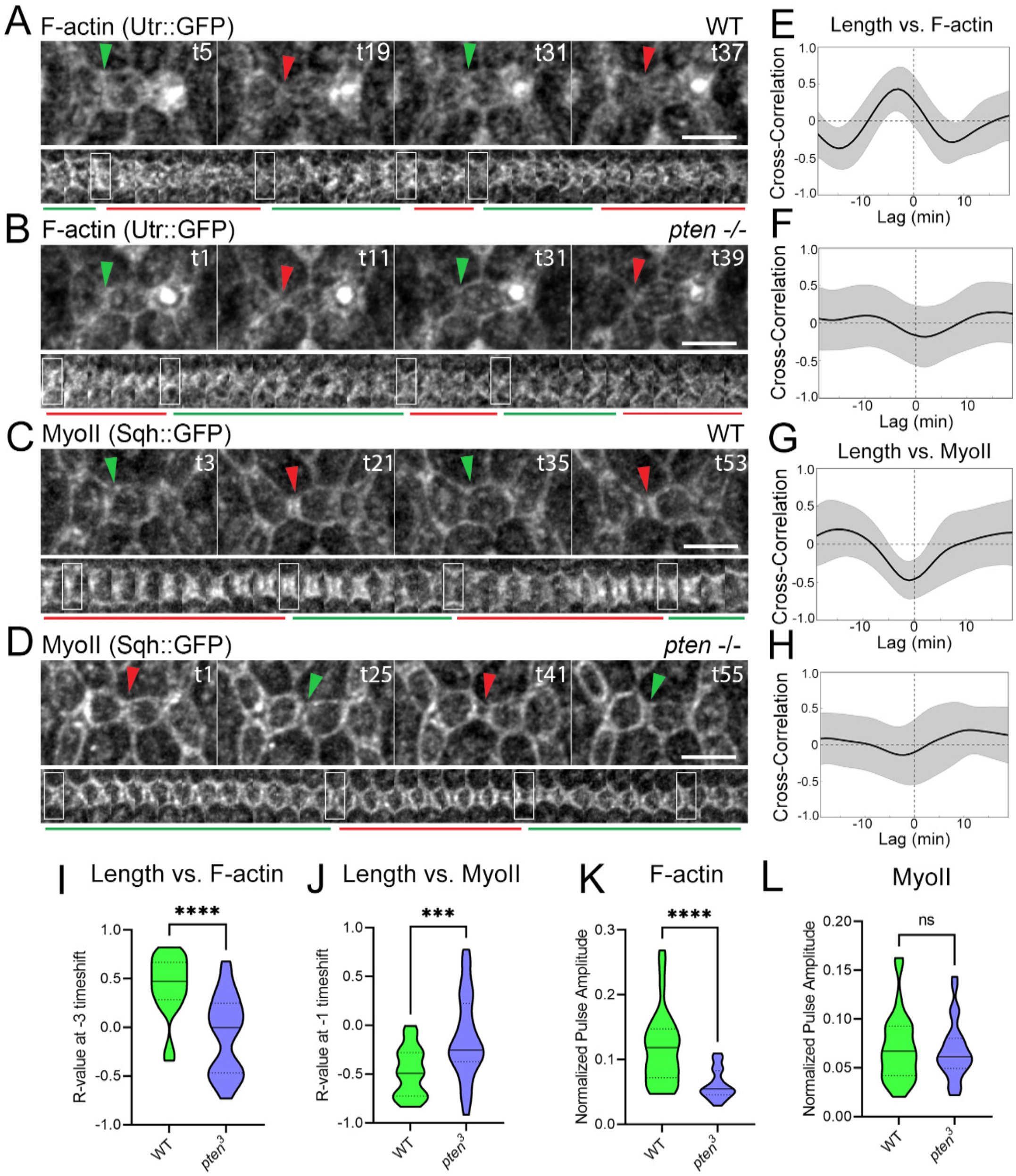
*pten* affects contractile and protrusive dynamics during contact pulsing. (A-D) Dynamics of (A-B) F-actin and (C-D) MyoII in (A, C) WT and (B, D) *pten* mutants, 26-28h APF. (E-H) Time-shifted Pearson’s correlations between contact length and levels of (E-F) F-actin and (G-H) MyoII in the above two genotypes. In this and subsequent graphs, the black line is the mean R-value and gray bar the SD. For each graph, n=30 contacts were pooled from 3 eyes. (E) Length vs. F-actin, WT. Peak R-value is .52 at a shift of −2. (F) Length vs F-actin, *pten*. Peak R-value is .13 at a shift of −7. (G) Length vs. MyoII, WT. Peak R-value is −.48 at a shift of −1. (H) Length vs MyoII, *pten*. Peak R-value is −.14 at a shift of-2. (I-J) Comparisons of correlations between length and (I) F-actin or (J) MyoII for individual WT and *pten* contacts at time-shift of WT peak R-value. n=30 contacts per genotype pooled from 3 eyes. (K-L) Comparison of pulse amplitudes of (K) F-actin and (L) MyoII normalized to reporter levels over 60 minutes at vertices of individual contacts. Statistics: Mann-Whitney *u* test, ***p<.001, ****p<.0001.

Additionally, the negative correlation between MyoII and contact length decreased as well (Fig. 2G-H, J). Moreover, the pulse amplitude of F-actin decreased, although the pulse amplitude of MyoII was not significantly different from WT (Fig. 2K-L). During periods of contraction, the parallel cables of MyoII that form along the LC-LC contacts were frequently further apart than in WT, suggesting that these contacts were either under greater tension, reduced protrusion, or both (Fig. 2C-D, red arrows). Because contractile and protrusive dynamics are mutually dependent in this system, disrupting one network may disrupt the other. The reduction in F-actin pulsing suggested a direct effect on actin branching and protrusive force. Overall, we find that *pten* affects both protrusive actin branching and contractile MyoII dynamics, supporting a role for *pten* in rebalancing the opposing forces generated by these networks.

### *pten* affects localization and dynamics of protrusive regulators at LC-LC contacts

The disruption of protrusive F-actin and contractile MyoII dynamics in *pten* mutant eyes may arise from disruption of the WRC, which regulates actin branching, or Rho1 GTPase signaling, which regulates actomyosin contractility, or both. The persistent accumulation of MyoII and the diminished pulsing of F-actin at LC-LC contacts suggested that F-actin assembly is affected. We therefore examined the dynamic accumulation of the WRC subunit Abi, which activates Arp2/3-mediated actin branching, at LC-LC contacts of *pten* mutant eyes compared with WT (Fig. 3A-B, Movie 3). To monitor WRC dynamics, we generated GFP-tagged Abi (GFP::Abi) controlled by the ubiquitin promoter (Akbari et al., 2009). We found strong pulse amplitudes of Abi at vertices during contact expansion in WT and significantly decreased pulse amplitudes and frequencies in *pten* mutant eyes (Fig. 3C-D). The time-shifted correlation between contact length and Abi levels at vertices also decreased in *pten* eyes compared with WT (Fig. 3E-G). These results highlight the role of the WRC at tAJs in controlling actin branching and protrusive force generation. They suggest that the defects in contact length pulsing, at least in part, arise from defects in WRC accumulation and actin branching at tAJs.

**Figure 3:**
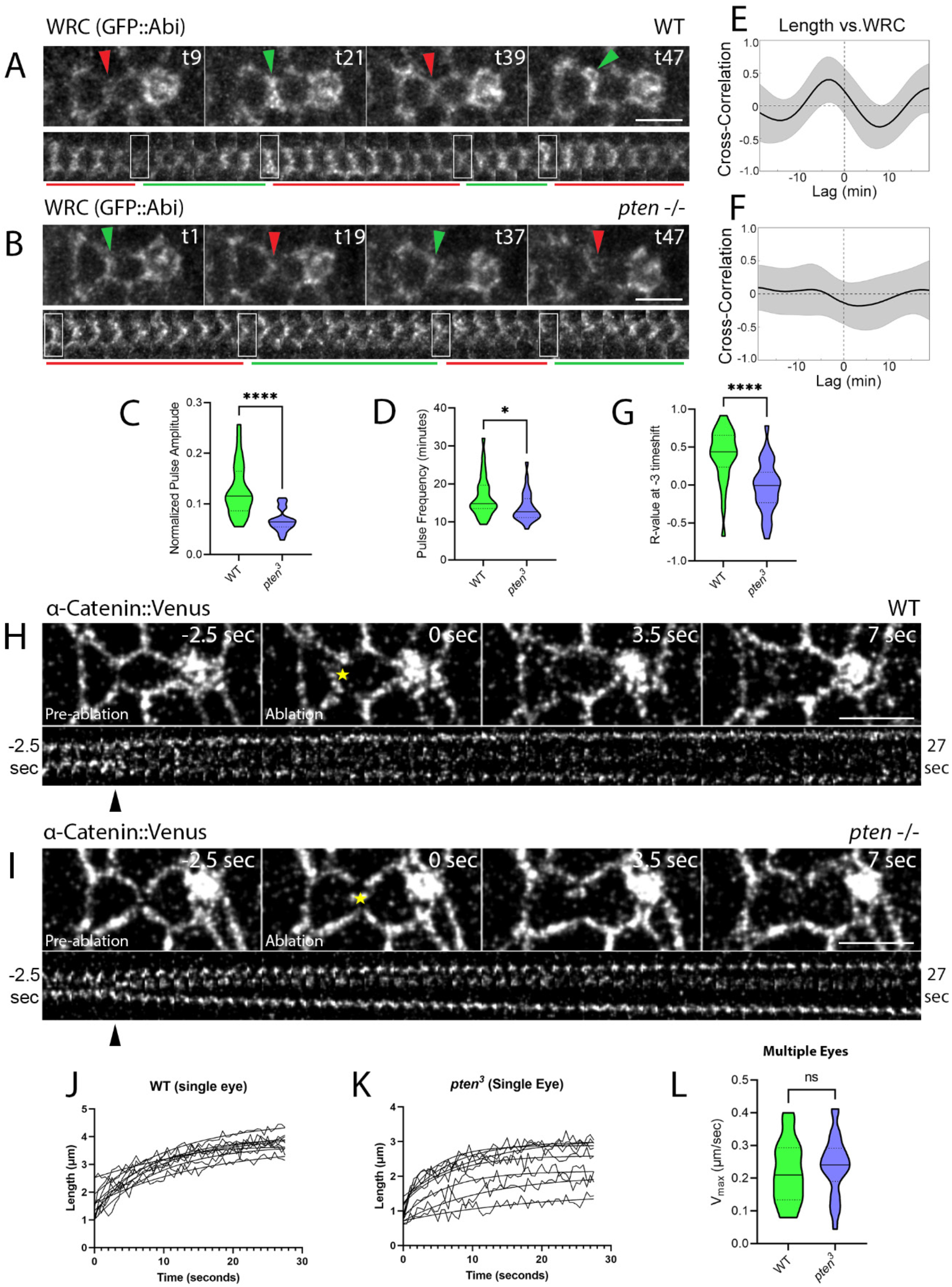
*pten* affects Abi dynamics during eye epithelial remodeling. (A-B) Dynamics of Abi in (A) WT eyes and (B) *pten* mutant eyes. (C) Comparison of pulse amplitude of Abi at vertices normalized to mean Abi. N=30 WT, 30 *pten* contacts pooled from 3-4 eyes per genotype. Unpaired t-test with Welch’s correction, p>.0001. (D) Comparison of Abi pulse frequency. Mann-Whitney *u* test, p=.0209. (E-F) Time-shifted correlations between contact length and Abi levels in WT and *pten* eyes. (E) WT. Peak R-value is .39 at a shift of −3. (F) *pten*. Peak R-value is .21 at a shift of −3. (G) Comparison of R-values at a shift of −3 in both genotypes. Unpaired t-test, p>.0001. (H-I) Snapshots and kymographs of laser ablations of LC-LC contacts in (H) a WT eye and (I) a *pten* eye. Yellow stars mark sites of ablation. (J-K) Recoil curves of 10 ablations each from (J) a single WT eye and (K) a single *pten* eye. (L) The initial recoil velocity (V_max_) of individual contacts pooled across 3 eyes in WT compared with *pten* mutants. Unpaired t-test, n=18-25 contacts.

The diminished amplitude of contact expansion may also arise from increased contraction. To distinguish between these possibilities, we measured the initial recoil velocities of vertices after ablating a small central region of LC-LC contacts in WT and *pten* mutant eyes as a proxy for tension along these contacts (Colombelli and Solon, 2013; Rauzi and Lenne, 2015) (Fig 3H-I). We found no difference in the recoil velocities between WT and *pten* mutant eyes, indicating that *pten* loss does not increase tension at LC-LC contacts (Fig. 3J-L). This result implies that Pten primarily promotes WRC activation and actin branching to support protrusive dynamics and contact expansion.

### Pten’s lipid phosphatase activity is required for epithelial remodeling

Because prior research implicated PIP_3_ in epithelial remodeling, and the *pten^c494^*allele eliminates Pten’s phosphatase activity, we hypothesized that Pten primarily affects the epithelium through its lipid phosphatase function (Bardet et al., 2013; Del Signore et al., 2018; Di Paolo and De Camilli, 2006; Huang et al., 1999). Point mutants of vertebrate PTEN (C124S and G129E) were previously shown to impair both its lipid and protein phosphatase activity or only its lipid phosphatase activity, respectively (Liliental et al., 2000; Myers et al., 1998; Qi et al., 2020). We introduced the equivalent point mutations (C132S and G137E, respectively) in *Drosophila* Pten to determine if Pten modulates PIP_3_ dynamics to control epithelial remodeling. We then used the MARCM technique to express each variant in *pten* mutant clones to determine if they could rescue the *pten* clonal phenotypes. We found that WT Pten rescued the *pten* clonal phenotypes, while *pten^C132S^* and *pten^G137E^*point mutants were unable to rescue these phenotypes (Fig. S2A-E, G). However, the clonal phenotypes were less severe in the *pten* clones expressing the *pten^G137E^* variant retaining protein phosphatase activity, suggesting that Pten’s protein phosphatase activity contributes to Pten function (Fig. S2C-D, G). Additionally, even the catalytically dead *pten^C132S^* variant accomplished minimal rescue, suggesting that Pten may have an additional non-catalytic function in the process (Fig. S2D, G). Together, these results indicate that Pten’s lipid phosphatase activity is essential for eye epithelial remodeling but suggest additional roles for other functions of Pten, including its protein phosphatase activity.

### Pten affects PIP_3_ dynamics

Since we found that Pten’s lipid phosphatase activity is required for proper epithelial remodeling, we examined Pten’s effects on PIP_3_ in developing eyes. We employed the PH domain of Grp1 fused to GFP (Grp1-PH::GFP or GPH) to monitor PIP_3_ in *pten* mutant clones and eyes (Britton et al., 2002). As expected, PIP_3_ levels were elevated in *pten* mutant clones compared with WT cells (Fig. 4A). To avoid confounding non-autonomous effects, we compared PIP_3_ dynamics in WT eyes with *pten* mutant eyes. Similar to WRC pulses, PIP_3_ pulses and contact expansion pulses co-occurred in WT eyes (Fig. 4B, Movie 4). In the *pten* mutant eyes, we found decreased pulse amplitude and increased pulse frequency of PIP_3_ (Fig. 4C-E, Movie 4). Moreover, the positive time-shifted correlation between PIP_3_ and contact length was abolished in *pten* eyes. In WT, PIP_3_ levels peaked 3 minutes before maximal contact expansion, while in *pten* mutants there was no such peak (Fig. 4F-I). Thus, PIP_3_’s predictable pulsed dynamics were eliminated in *pten* mutants and remained mostly static.

**Figure 4:**
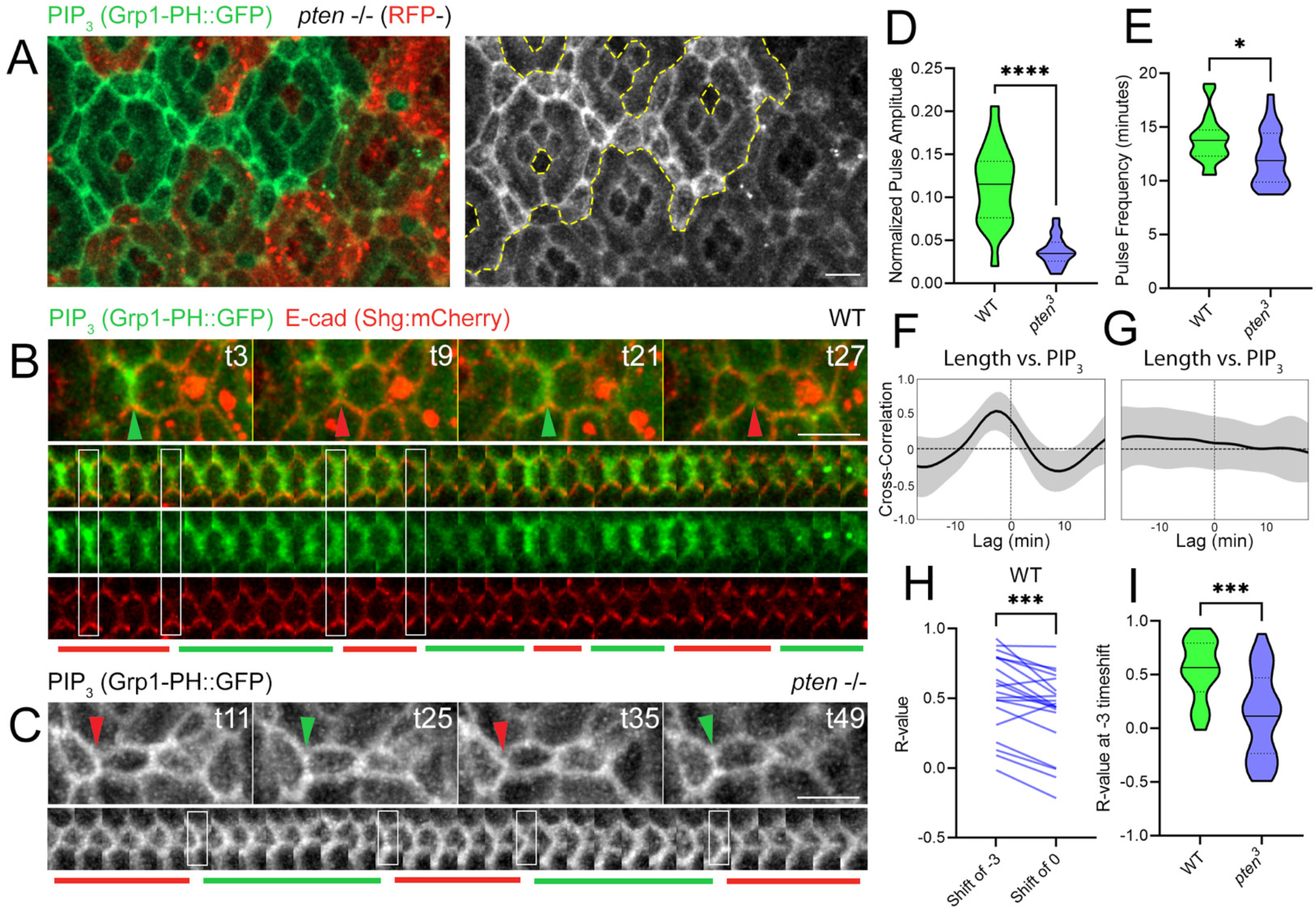
*pten* affects PIP_3_ dynamics during eye epithelial remodeling. (A) Levels and localization of PIP_3_ in *pten* mutant clones, 28-30h APF. (B-C) PIP_3_ dynamics in (B) WT eyes and (C) *pten* mutant eyes. (D) Comparison of pulse amplitude of PIP_3_ normalized to mean PIP_3_. N=20 WT, 29 *pten* contacts pooled from 2-3 eyes per genotype. Unpaired t-test with Welch’s correction, p<.0001. (E) Comparison of PIP_3_ pulse frequency. Unpaired t-test, p=.04975. (F-G) Time-shifted correlations between contact length and PIP_3_ levels in WT and *pten* eyes. (F) WT. Peak R-value is .54 at a shift of −3. (G) *pten*. Peak R-value is .02 at a shift of +13. (H) Comparisons of R-values at a shift of −3 vs. a shift of 0 in WT. Paired t-test, p=.0002. (I) Comparison of R-values at a shift of −3 in both genotypes. Unpaired t-test, p=.0002.

### Pi3K affects eye epithelial remodeling

To further determine the role of PIP_3_ dynamics in epithelial remodeling, we examined the role of Pi3K, the enzyme that phosphorylates PIP_2_ to PIP_3_. We asked whether increasing Pi3K activity could mimic the *pten* mutant phenotype and if loss or downregulation of Pi3K function affects epithelial remodeling. Active Pi3K consists of a catalytic subunit, Pi3K92E (Pi3K-cat), and a regulatory subunit, Pi3K21B (Pi3K-reg) (Leevers et al., 1996; Weinkove et al., 1997). To increase Pi3K activity, we expressed a constitutively active, membrane-tethered Pi3K-cat (PI3K-cat.CAAX) (Leevers et al., 1996). To decrease *Pi3K* activity, we used a *Pi3K-reg* deletion mutant generated using gene editing and eyes and clones depleted of Pi3K-cat using RNA-mediated interference (RNAi) (Trivedi et al., 2020). We confirmed the increase or decrease of Pi3K function by monitoring PIP_3_ levels with GPH in Pi3K-cat.CAAX- and Pi3K-cat RNAi-expressing clones and *Pi3K-reg* mutant clones (Fig. S3A-C, K-L). We found that increasing Pi3K-cat function led to pinched contacts and defects similar to those in *pten* mutant eyes, such as rosettes and clustered bristles. However, the aberrant 1°-1° contacts common in *pten* mutant eyes were rare (Fig. S3A, D, K). Depleting Pi3K-cat function also caused epithelial remodeling defects, with some similar to those in *pten* mutants, including rosettes and aberrant 1°-1° contacts (Fig. S3E, L). Similar defects were also found in *Pi3K-reg* mutant eyes (Fig. S3F).

As both the increase and decrease of Pi3K and PIP_3_ levels caused defects in cytoskeletal regulation, we examined PIP_3_ dynamics in clones with increased and decreased Pi3K activity. We found that PIP_3_ pulsing and correlation with contact length were reduced in Pi3K-cat.CAAX clones (Fig. 5A-B, Movie 5). In Pi3K-depleted clones, PIP_3_ fluctuations appeared reduced as well (Fig. S3J, Movie 5). We additionally measured the amplitude of contact length pulsing in these clones and in *PI3K-reg* mutant eyes. Contact length amplitude was reduced in *PI3K-reg* mutants compared to WT and showed a trend toward reduction in both PI3K-cat.CAAX- and PI3K-cat RNAi-expressing clones (Fig. 5C-D, Fig. S3K-N).

**Figure 5:**
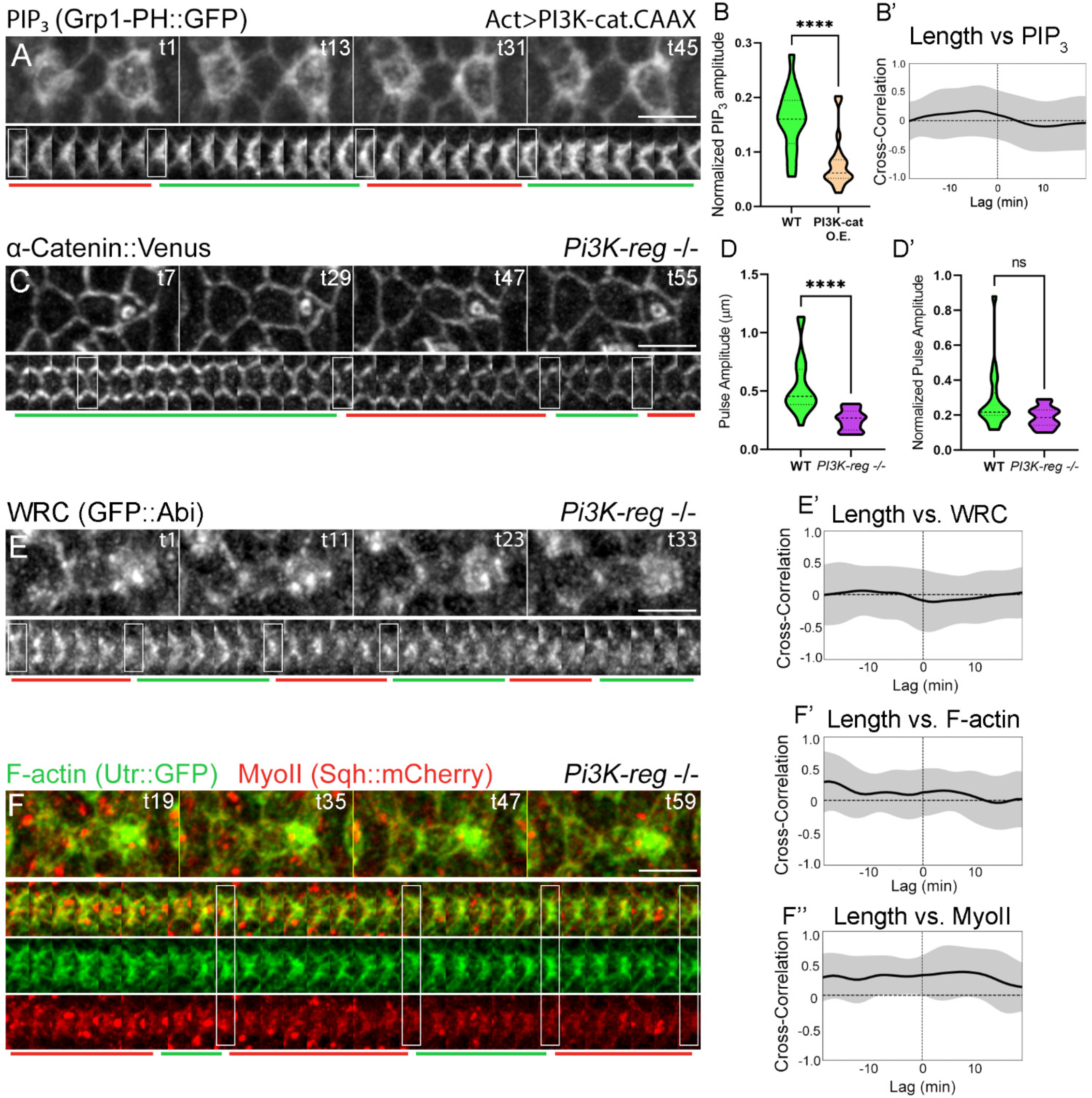
Pi3K Loss- and gain-of-function disrupt eye epithelial remodeling. (A) PIP_3_ pulsing in clones expressing Pi3K-cat.CAAX (B) PIP_3_ pulse amplitudes normalized to total PIP_3_ in clones compared with WT regions. Mann-Whitney *u* test, p<.0001, n=16-22 contacts pooled from 4 eyes. (B’) Time-shifted correlations between contact length and PIP_3_ in clone regions expressing Pi3K-cat.CAAX (peak correlation .17 at a shift of −5). (C) LC-LC contact dynamics in *Pi3K-reg* mutant eyes. (D) Quantification of (D) absolute and (D’) normalized pulse amplitudes in *Pi3K-reg* eyes compared with WT. Mann-Whitney *u*-test, n=20 contacts ****p<.0001. (E-F) Dynamics of (E) the WRC and (F) F-actin and MyoII in *Pi3K-reg* eyes. (E’, F’) Time-shifted correlations between contact length and (H’) the WRC (peak correlation −.11 at shift of +2), (F’) F-actin (peak correlation .15 at a shift of +4), and (F’’) MyoII (peak correlation .30 at a shift of −3).

To further characterize the *Pi3K* loss-of-function phenotype, we examined F-actin, MyoII, and WRC dynamics in *Pi3K-reg* mutant eyes (Movie 6). Similar to *pten* mutant eyes, we found decreased correlations between contact length and the WRC (Fig 5E), F-actin and MyoII (Fig. 5F). Finally, to further test Pten’s relationship with Pi3K, we generated *pten* mutant clones expressing Pi3K-cat RNAi. Depleting *Pi3K-cat* function partially rescued the *pten* phenotype, further indicating a role for PIP_3_ in this process (Fig. S2F-G). These results demonstrate that Pi3K regulates force generation and transmission and suggest that PIP_3_ cycling, in addition to overall levels, is required for proper epithelial remodeling.

### Pten affects Rac1 function while Rac1 affects PIP_3_ and protrusive dynamics

Because *pten* mutants disrupt PIP_3_, the WRC, and protrusive dynamics and because Rac1 and PIP_3_ are known to activate the WRC *in vitro*, we tested Rac1 function in this process (Fig. 6A-B, Movie 7). We found that the amplitude and frequency of Rac1 fluctuations were reduced in *pten* mutants compared with WT, indicating impaired dynamics (Fig. 6C-D). Additionally, we found a slight but significant positive correlation between Rac1 and LC-LC contact length in WT. This correlation was abolished in *pten* mutants (Fig. 6E-G).

**Figure 6:**
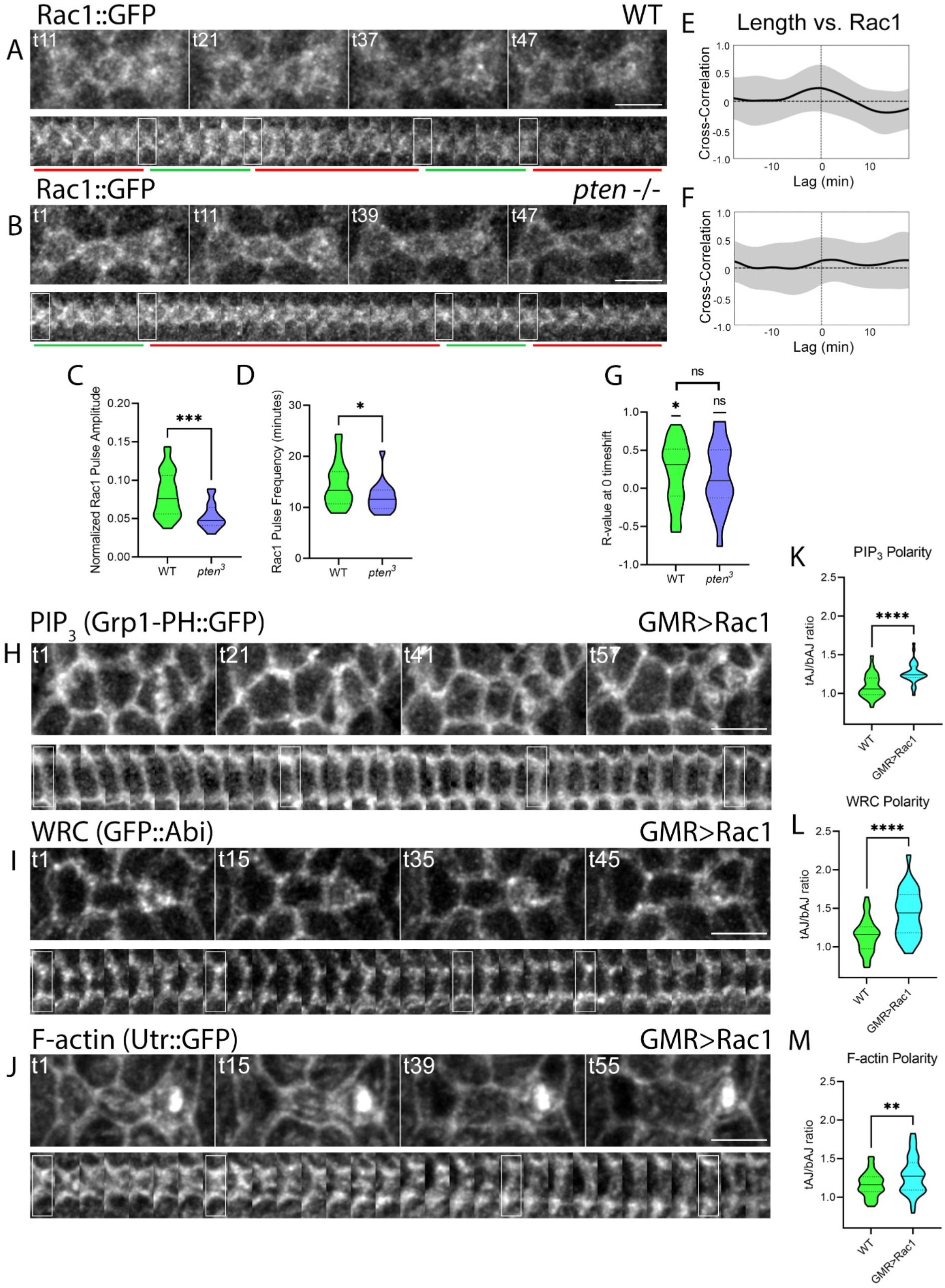
*pten* affects Rac1 signaling, which affects the WRC and protrusive F-actin dynamics. (A-B) Rac1::GFP dynamics in (A) a WT and (B) a *pten* mutant eye. (C-D) Comparisons of pulse (C) amplitudes and (D) frequencies of Rac1 at contact vertices in the two genotypes. (T-test, n=28 WT, 20 *pten*, *p<.05, ***p<.001). (E-F) Time- shifted correlations between contact length and Rac1 levels in WT and *pten*. (E) WT: peak R-value = .21 at a shift of 0, significantly different from R-value of 0, one-sample t-test, p=.0108, n=28 contacts from 3 eyes (F) *pten*: peak R-value =.16 at a shift of +1, no significant difference from R-value of 0, one sample t-test, p=.1493, n=20 contacts from 2 eyes. (G) Comparison between R-values at 0 for both genotypes, unpaired t-test, p=.6800. (H-J) Dynamics of (H) PIP_3_ (I) the WRC and (J) F-actin in eyes overexpressing Rac1. Ordinary patterns of accumulation along with contact expansion and contraction are abolished. (K-M) Quantifications of localization of (K) PIP_3,_ (L) the WRC and (M) F-actin in eyes overexpressing Rac1. All three molecules are more concentrated at vertices in Rac1-overexpressing eyes compared with control. Unpaired t-tests, **p<.01, ****p<.0001. n=29-66 contacts pooled from 2-4 eyes per genotype.

It was previously shown that *rac1*, *rac2* mutant retinae have defects in epithelial organization (Martin-Bermudo et al., 2015). Considering this role of *rac* genes and their potential interaction with *pten*, we overexpressed Rac1 and investigated its effects on epithelial remodeling. Broad Rac1 expression resulted in epithelial remodeling defects and altered mechanical and cytoskeletal dynamics. The apical area of epithelial cells was enlarged and cells dynamically expanded and relaxed, in contrast to the static, pinched LC-LC contacts observed in *pten* eyes. While in WT, PIP_3_, Abi, and F-actin are dynamically enriched at vertices and LC-LC contacts, in Rac1 expressing eyes they concentrated more strongly at vertices and additionally localized along the entire cell perimeter at the level of AJs. Moreover, PIP_3_ accumulated in pulses along the apical cell area, implying that Rac1 activates Pi3K and PIP_3_ production. Together, these effects suggest that Rac1 and PIP_3_ activate the WRC and protrusive F-actin dynamics in part through the activation of Pi3K in a positive feedback mechanism (Fig. 6H-J, Movie 8). The overexpansion of LCs often led to tears in the epithelium that coincided with waves of PIP_3_, Abi and F-actin along the rupturing cell perimeter (Fig. S4A-C, Movie 9).

Clonal overexpression of Rac1 induced similar epithelial tears and fused cells (Fig. S3D). Overexpressing Rac1 in *SCAR* mutant clones decreased the number of ruptured cells, with fewer than half the eyes observed exhibiting this phenotype likely due to residual *SCAR* function in the clones (Fig. S3E). Rac1 overexpression in *pten* clones failed to induce ruptures (Fig. S3F). These results suggest that Rac1 acts through the WRC to promote protrusive F-actin dynamics and apical area expansion and requires proper PIP_3_ dynamics to function.

Overall, our findings show that Pi3K dynamics promote Rac1 function, which in turn influences WRC activation, actin branching, and protrusive force generation. In addition to activating the WRC and protrusive dynamics indirectly through Rac1, PIP_3_ dynamics can activate the WRC directly and thus cooperate with Rac1 to maximally activate the WRC. Rac1 also appears to sustain and amplify the PIP_3_ signal, to promote actin branching and pulsatile LC-LC contact expansion.

### Pten and Pi3K localize dynamically in the vicinity of LC-LC contacts

If Pi3K and Pten directly affect LC-LC contact pulsing, they would likely localize to or near these contacts and associated tAJs. Pi3K-reg is primarily responsible for the spatiotemporal activation of Pi3K (Fruman, 2010). Therefore, to identify the sites of Pi3K activation, we generated a GFP-tagged Pi3K-reg (Pi3K-reg::GFP) driven by the ubiquitin promoter. We found that Pi3K-reg::GFP accumulated dynamically at the level of AJs. During early expansion, Pi3K-reg::GFP localized to four spots on either side of tAJs. As the contacts expanded, the domain of Pi3K-reg localization expanded and moved toward the tAJs (Fig. 7A, Movie 10). To quantify this, we measured the temporal correlation between Pi3K at or adjacent to vertices and overall contact length and found that the adjacent-to-vertex correlation peaked earlier than the at-vertex correlation (Fig. 7B). Pi3K both at and adjacent to the vertex correlated strongly with F-actin (Fig. 7C). Therefore, Pi3K localization around tAJs is po mediate LC-LC contact expansion by promoting protrusive dynamics and pushing the plasma membrane outward. To further investigate this process, we used immunostaining to examine Pi3K-reg::GFP localization relative to E-cad and Sdk, which marks tAJs (Malin et al., 2022). When Pi3K-reg localized in spots at a distance from tAJs, it colocalized strongly with E-cad. In addition, Pi3K-reg localized with or partially overlapped with Sdk at tAJs (Fig. 7D).

**Figure 7:**
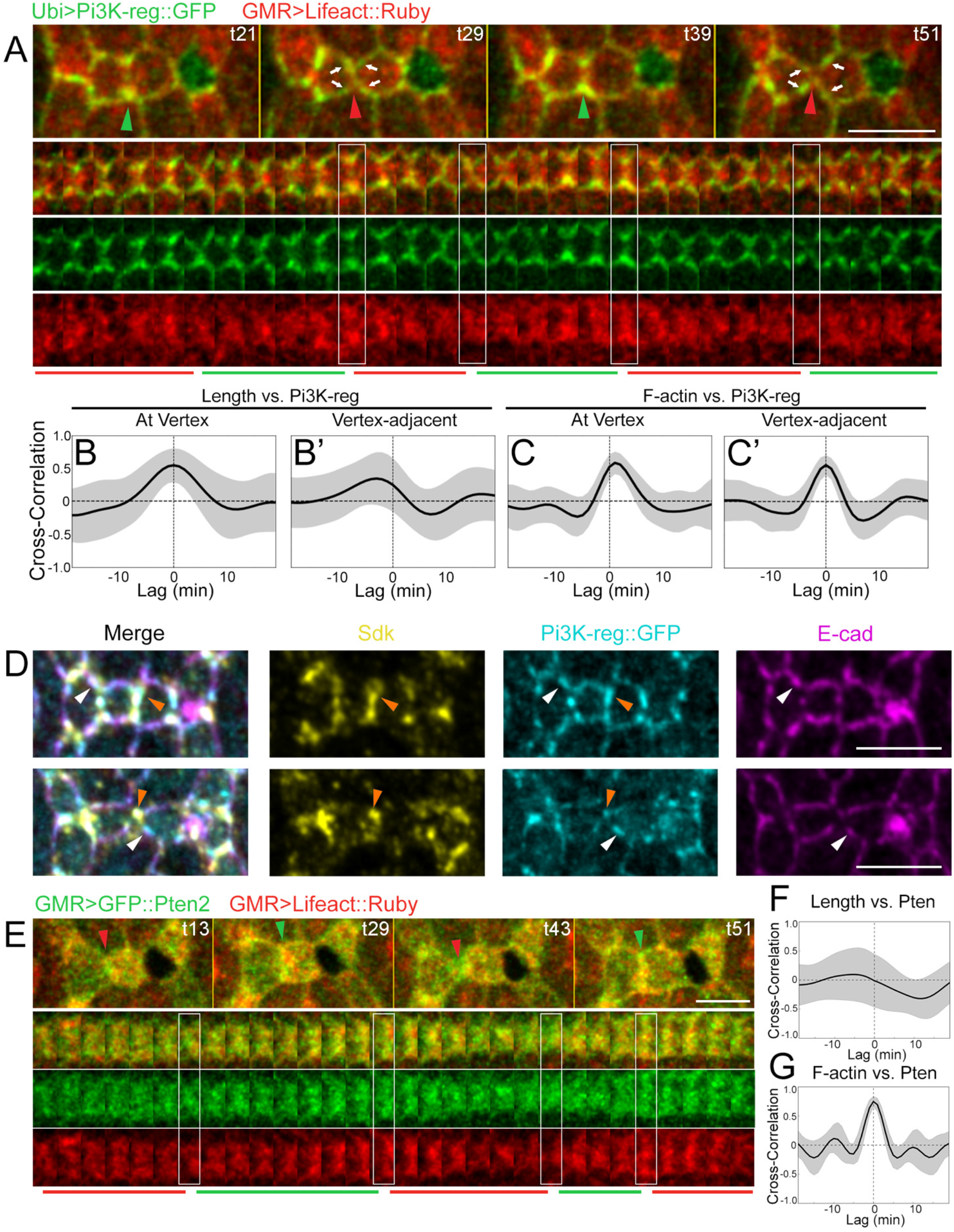
Pi3K-reg accumulates preferentially and dynamically along LC-LC contacts. (A) Pi3K-reg::GFP and F-actin dynamics in a WT eye. Pi3K-reg localizes to vertices in expanded contacts, and in spots around the vertex in contracted contacts (white arrows). (B) Time-shifted correlations between contact length and Pi3K- reg (B) directly at the vertex (peak R-value .54 at a shift of 0) and (B’) adjacent to the vertex, (peak R-value .34 at a shift of −3). (C) Time-shifted correlations between F-actin and Pi3K-reg (C) at the vertex (peak R-value=.58 at a shift of +1) and (C’) adjacent to the vertex (peak R-value=.55 at a shift of 0). N=30 contacts pooled from 3 eyes for each correlation. (D) Two rows of cells expressing Pi3K-reg::GFP and stained for Sdk and E-cad. White arrowheads: colocalization between Pi3K and E-cad. Orange arrowheads: colocalization between Pi3K and Sdk. (E) GFP::Pten and F-actin dynamics in a WT eye. (F-G) Time correlations of Pten levels with (F) contact length and (G) F-actin. Pten does not show a strong correlation with contact length (peak R-value .10 at shift of −9) but does correlate with F-actin (peak R-value .67 at a shift of 0).

Pten activation requires membrane localization through its PDZ domain binding motif and phospholipid binding (Das et al., 2003; Jang et al., 2021). We created a GFP::Pten reporter driven by the ubiquitin promoter to identify the membrane sites where Pten counteracts Pi3K. We tagged the Pten2 isoform, which localizes to apical junctions (Maehama et al., 2004; Pickering et al., 2013; von Stein et al., 2005). We found that Pten accumulated in a diffuse pattern both at and surrounding LC-LC contacts (Fig. 8E, Movie 12). Although Pten localization was dynamic and correlated with F-actin levels, Pten distribution was relatively uniform during both contact expansion and contraction (Fig. 8F-G). These results suggest that changes in PIP_3_ levels are caused primarily by fluctuations in Pi3K levels around tAJs, while PIP_3_ turnover requires both Pi3K and Pten.

## DISCUSSION

After the initial cellular organization of ommatidia through the cell-specific distribution of adhesion molecules, continuous cell movement is required for the precise geometry of the eye to develop. Regular pulses of contraction and expansion of cell contacts are characteristic of this movement. Here, we show that Pten and Pi3K regulate oscillating levels of PIP_3_ at apical junctions and this is necessary for normal pulsing shape changes and morphogenesis. The function of PIP_3_ is to generate protrusive force and elongate contacts by activating the WRC and thereby promoting actin branching. The dependence of protrusion on dynamic changes in PIP_3_ combined with the proximity of Pi3K and Pten to each other near junctions makes this system uniquely suited to precise temporal control of protrusion. We provide evidence that phosphoinositides controlled by Pten and Pi3K affect Rac1 function and propose that PIP_3_ cooperates with Rac1 to activate the WRC and protrusive actin branching to control cytoskeletal and contact length pulsing.

### Technical considerations

While phosphoinositides are widely studied, little of this research relates to planar epithelial morphogenesis. In addition to looking at cell shape, we utilized live imaging of cytoskeletal indicators of both contractile and protrusive forces, as the control of protrusion in planar epithelial development remains largely unexplored. Dynamic imaging of PIP_3_ allowed us to quantify its cyclical ebb and flow in addition to simple levels. We used engineered point mutants to differentiate between the lipid phosphatase and protein phosphatase functions of Pten. In addition, we directly investigated Pi3K function using a novel null mutation generated by gene editing (Trivedi et al., 2020) and verified it by loss of PIP_3_ and the associated phenotype. Further, by generating a *Pi3K-reg* mutant clones, we showed the feasibility of studying Pi3K-null clones. This contrasts with previous work relying on dominant-negative suppression of Pi3K function in which residual function or unanticipated effects remain a consideration. Genetic tests for expressing Rac1 in *SCAR* and *pten* mutant clones imply that Pten and SCAR are important for the function of Rac1. Taken together these approaches provide new insight into the role of Pten, Pi3K, and PIP_3_ in regulating protrusive forces and planar epithelial morphogenesis.

Our results show that in *pten* mutants cell contacts are pinched and fail to lengthen, a result that echoes an earlier study of notum development (Bardet et al., 2013). In that case, the mechanism of this effect was linked to persistent and elevated contractility. However, since that time, the importance of protrusive forces in contact remodeling has been demonstrated, including strong pulsatile PIP_3_ accumulation coinciding with the accumulation of the WRC and protrusive F-actin (Del Signore et al., 2018; Malin et al., 2022; Patel et al., 2008). This observation aligns with literature linking PIP_3_ with WRC activation in other contexts (Chen et al., 2017; Hume et al.; Koronakis et al., 2011; Mendoza, 2013; Mendoza et al., 2011; Schaks et al., 2018). Taken together they suggest that the *pten* phenotype is caused by dysregulation of protrusive forces, even though the contacts are shortened. Key results supporting this idea in our system are that the WRC does not accumulate in *pten* mutants, there is no evidence for changes in MyoII levels and, although contacts are pinched, when laser ablated their initial recoil velocity does not change, indicating no change in contractile forces. Therefore, our results put Pten’s function in planar epithelial morphogenesis in a new light, establishing its role in regulating protrusive forces in this context.

Because Pten can act through many intermediaries and has both lipid and protein phosphatase activity, investigating its mechanism of action is important. Our results confirm the importance of PIP_3_ in this system. Specifically, we show that a *pten* point mutant with a deficient lipid phosphatase function fails to rescue the *pten* phenotype. In addition, we examine the role of Pi3K, which opposes Pten in the cycle of phosphorylation and dephosphorylation that regulates PIP_3_ levels. Mutations affecting Pi3K also show epithelial remodeling defects. Moreover, we show a partial rescue of the *pten* mutant phenotype by perturbing Pi3K’s catalytic subunit. Though our results leave open the possibility of other effects through Pten’s protein phosphatase activity, they establish the importance of PIP_3_ regulation by Pten and Pi3K in the observed phenotypes.

Strikingly, we found that genetic manipulations that either increased or decreased levels of PIP_3_ resulted in similar phenotypes. In phenotypes in which PIP_3_ levels were constitutively low due to reduced Pi3K activity, contacts were shortened due to a deficit in protrusion. However, with the elevated PIP_3_ levels in *pten* mutants or with constitutively active Pi3K, LC-LC contacts were also short or lost, with a decrease of WRC accumulation and consequently branched actin. This is opposite to what would be expected given the ability of PIP_3_ to activate the WRC and protrusive dynamics (Chen et al., 2010; Hume et al.; Lebensohn and Kirschner, 2009; Oikawa et al., 2004; Suetsugu et al., 2006). It has been noted in other contexts that there exists continuous turnover of various phosphoinositides and it remains an open question if this turnover is integral to their mechanism of action (Di Paolo and De Camilli, 2006). The current results reveal that high levels of PIP_3_ alone are not sufficient for marshaling protrusive forces in this system, and that dynamism of PIP_3_ is also critical. Speculatively, this dependence could optimize the sensitivity of the system and its ability to rebalance mechanical forces, as turnover would allow the balance of different phosphoinositides to rapidly shift. Alternatively, the mechanisms needed for proteins to continuously regenerate their activity in the ongoing process of remodeling actin networks may intrinsically require phosphoinositide levels to fluctuate.

In contracted contacts under high tension, Pi3K-reg localizes at four spots at some distance from the tAJs, and PIP_3_ levels are low, indicating low Pi3K lipid-kinase activity. During expansion, Pi3K-reg’s localization spreads away from these spots, concentrating at tAJs as PIP_3_ levels peak. Thus, Pi3K-reg’s distribution and its lipid kinase function change with tension. Pten, in contrast, does not detectably shift distribution through the pulse cycle, but its levels increase with increasing F-actin. The specific mechanisms that regulate Pi3K and Pten distribution here are unknown, but the proteins are clearly correlating with oscillations in mechanical forces. Likewise, during pulsing, Sdk concentrates at tAJs in response to tension and associates with the WRC to reduce tension and promote contact expansion. Pyd, on the other hand, accumulates following maximal contact expansion to increase tension and promote contraction (Malin et al., 2022). Pi3K and Pten join Sdk and Pyd as elements that change in response to pulsing and are moreover molecular effectors of mechanical forces that generate pulsing. Thus, Pi3K and Pten appear as integral components to the feedback loops that generate and maintain pulsing shape changes.

Although they exhibit different patterns of distribution that vary with the pulse cycle, both Pi3K and Pten are enriched at and near a subset of cell contacts at the level of tAJs. The spatial proximity between the two proteins is consistent with their capacity to temporally control PIP_3._ This is an interesting contrast with other systems in which Pi3K and Pten do not colocalize but are rather separated to define distinct subcellular domains. During epithelialization, Pten localizes apically, while Pi3K localizes basolaterally, to define the apicobasal axis (Martin-Belmonte et al., 2007; Pinal et al., 2006; von Stein et al., 2005). In motile cells, Pi3K localizes at the lamellipodium while Pten is at the rear and sides to coordinate protrusions with contractions (Funamoto et al., 2002; Iijima et al., 2002). Thus, the proximity of Pi3K and Pten at apical junctions positions them for precise temporal regulation of PIP_3_ pulsatile dynamics in this location.

Our data support the idea that PIP_3_ regulates the WRC both directly and at least in part indirectly through Rac1 activation. Loss of *pten* reduces the pulsing of Rac1 levels, suggesting that PIP_3_ activates Rac1. This would be consistent with the documented roles of PIP_3_ and Rac1 in promoting the formation of lamellipodia and with current models of WRC activation (Chen et al., 2017; Chen et al., 2010; Ridley et al., 1992). We also found that overexpressing Rac1 induced protrusive F-actin and PIP_3_ accumulation causing expansion and rupture of cells’ apical area. This is in contrast to manipulations that raised PIP_3_ levels directly that did not increase protrusion. Together these data suggest that PIP_3_ could activate both a GEF for Rac1 at apical junctions and that Rac1 itself might then activate Pi3K to sustain its activation and generate a pulse of WRC activation and actin branching (Kovacs et al., 2002; Nakagawa et al., 2001; Perez et al., 2008). Thus, Rac1 could act not only as an effector of PIP_3_ in activating the WRC, but also a feedback amplifier of PIP_3_ signaling and through these actions together would have the capacity to robustly promote actin branching at apical junctions.

In sum, we show that Pten and Pi3K affect PIP_3_ pulsing to control cytoskeletal and mechanical pulsing. Characterization of multiple genetic conditions indicates that dynamic changes in PIP_3_ levels are essential for promoting protrusion. PIP_3_ pulsing activates the WRC and actin branching, and this system utilizes Rac1 to promote protrusive dynamics and reinforce PIP_3_ production through positive feedback. Additionally, Pten levels and both the distribution and the activity of Pi3K are regulated with cycles of contact expansion and contraction, suggesting regulation by mechanical signals. Together these results reveal the importance of Pten, Pi3K and PIP_3_ in generating protrusive forces at tAJs to control planar epithelial morphogenesis.

## ACKNOWLEDGEMENTS

We thank E, Hafen, Daniel St Johnston, T. Millard, R. Padinjat, F. Pichaud, A. Martin, J. Treisman and J. Zallen for generous gifts of fly strains, and J. Treisman for antibodies, the Bloomington Stock Center, the Vienna *Drosophila* Research Center, and the Kyoto Stock Center for fly strains, the Developmental Studies Hybridoma Bank for antibodies, and G. Rong and the Institute for Chemical Imaging of Living System (CILS) at Northeastern University for assistance with multi-photon imaging. We thank K. G. Commons for her critical reading of the manuscript and editorial suggestions. This work was supported by a grant from the National Institute of Health to V.H. (R01 GM129151).

## Author Contribution

Conceptualization, V. Hatini; Investigation and Resources, J. Malin, C. Rosa and V. Hatini; Formal Analysis, J. Malin; Writing Original Draft, V. Hatini; Manuscript Review and Editing, J. Malin, C. Rosa and V. Hatini; Supervision, V. Hatini; Funding Acquisition, V. Hatini.

## Data availability

All data needed to evaluate the conclusions in the paper are present in the article and/or the supplementary materials. All image data will be deposited in the Tufts Metaverse database and given an accession number prior to publication. Image data and other supporting data of this study are available from the corresponding author upon reasonable request. Requests for reagents should be directed to and will be fulfilled by the lead contact, Victor Hatini.

## METHODS

### Fly strains

To examine the *pten* loss-of-function phenotypes, we employed the *pten^c494^*, *pten^3^*, and *pten^2L100^*alleles recombined to an FRT40A site to generate genetically marked mutant clones and fully mutant eyes. We used *pten^C494/3/2L100^* FRT40A; α-Catenin::Venus^CPTI^ (α-Cat::Venus) to examine *pten*’s role in epithelial rearrangements and *pten^3^*FRT40A with cytoskeletal reporters on chromosome III to examine *pten*’s role in cytoskeletal control. To determine if Pten’s lipid phosphatase or protein phosphatase activity affects epithelial remodeling, we constructed UAS-GFP::Pten2 (C132S), a point mutant that disrupts both the protein phosphatase and lipid phosphatase activities, UAS-GFP::Pten2 (G137E), a point mutant that preferentially disrupts the lipid phosphatase activity, and WT UAS-GFP::Pten2. We selected UAS insertions on the third chromosome with similar expression levels to determine if these variants can rescue the *pten* clonal phenotypes using the MARCM technique. To examine Pten and Pi3K-reg protein distribution and dynamics, we employed UAS-GFP::Pten2 and constructed a Ubi-GFP::Pten2 and Ubi-Pi3K-reg::GFP. Additionally, we constructed Ubi-GFP::Abi to track the dynamics of the WRC.

Fly lines from the Bloomington *Drosophila* Stock Center: (1) UAS-Lifeact::Ruby, (2) *sqh^AX3^*/FM7c; Sqh-Sqh::GFP, (3) tGPH; *Sb/TM3*, (4) tGPH (III), (5) Rac1-GFP::Rac1 (III), (6) *y w*, UAS-Pi3K92E.CAAX, (7) UAS-Pi3K92E RNAi (III), (8) UAS-Rac1 (II), (9) UAS-Rac1 (III), (10) GMR-GAL4 (III), (11), Ey-GAL4 (II), (12) UAS-dicer (II), (13), UAS-dicer (III), (14) hsFLP (II), (15) Ey-FLP (II), (16) GMR-Hid FRT40A/Cyo-GFP; Ey-GAL4 UAS-FLP, (17) *w* Ey-FLP; GMR-myrRFP FRT40A, (18) *SCAR*^𝛥^*^37^* FRT40A, and (19) pw+ 21C, pw+ 36F, FRT40A. α-Cat::Venus^CPTI002596^ was obtained from the Kyoto stock center.

Additional stocks used: (1) Sqh-UtrABD::GFP (III),(2) *sqh^AX3^*; sqh-UtrABD::GFP, Sqh-Sqh::mCherry (II), (3) Sp/CyO; Sqh-UtrAbd::GFP, Sqh-Sqh::mCherry/MKRS, and (4) Sqh-UtrABD::mCherry (II) (gifts of A. Martin), (5) UAS-Lifeact::Ruby; GMR-GAL4 (previously described by Del Signore et al.), (6) Ubi-Ecad::GFP, (7) Shg-Shg::mCherry ; tGPH, (8) Ey-GAL4 ; UAS-dicer (9) UAS-dicer ; GMR-GAL4.

The following stocks were gifts as indicated: Ey-FLP, UAS-GFP; tub-GAL80 FRT40A; Ubi-GAL4 (J. Treisman), *pten^3^* FRT40A and *pten^2L100^* FRT40A (E. Hafen), *pten^c494^* FRT 40A (D. St Johnston), *Pi3K21B^!−.^* indels (2,9) (P. Raghu). A *Pi3K21B^!−.2,9^* FRT40A recombinant was created using indel (2,9).

Fly lines that were generated in this study: (1) *pten^3^* FRT40A; α-Cat::Venus, (2) *pten^2L100^* FRT40A ; α-Cat::Venus, (3) *pten^c494^* FRT40A; α-Cat::Venus, (4) *pten^3^* FRT40A; Ubi-Abi::GFP, (5) pten^3^ FRT 40A; Sqh-Sqh::GFP, (6) *pten^3^* FRT40A; Sqh-Utr::GFP, (7) *pten^3^* FRT40A; tGPH, (8) *pten^3^* FRT40A; Rac1-GFP::Rac1 (BL-52285), (9) *pten^3^* FRT40A; UAS-GFP::Pten2, (10) *pten^3^* FRT40A; UAS-GFP::Pten2^C132S^, (11) *pten^3^* FRT40A; UAS-GFP::Pten2^G137E^, (12) *pten^3^* FRT40A ; UAS-Pi3K92E RNAi, (13) FRT40A ; UAS-GFP::Pten2, (14) *Pi3K21B^!−.2,9^* FRT40A, (15) *Pi3K21B^!−.2,9^* FRT40A ; a-Cat::Venus, (16) *Pi3K21B^!−.2,9^* FRT40A ; Sqh-Utr::GFP, Sqh-Sqh::mCherry, (17) *Pi3K21B^!−.2,9^* FRT40A ; Ubi-GFP::Abi, (18) *Pi3K21B^!−.2,9^* FRT40A ; tGPH, (19) *y w*, UAS-Pi3K92E.CAAX (BL-8294) ; Ey-FLP, (20) hsFLP ; UAS-Pi3K92E.RNAi (BL-27690), (21) ubi-Ecad::GFP ; Act>>GAL4, UAS-RFP, (22) tGPH (BL-8163) ; Act>>GAL4 UAS-RFP, (23) UAS-Rac1 (BL-58816) ; Sqh-Utr::GFP, Sqh-Sqh::mCherry, (24) tGPH ; UAS- Rac1 (25) Sqh-UtrABD::mCh ; GMR-GAL4, ubi-GFP::Abi, (26) SCAR*^!−.37^* FRT40A; UAS-Rac1 (BL-28874), (27) *pten^3^* FRT40A; UAS-Rac1, (28) FRT 40A; UAS-Rac1.

### Genetic analysis

GMR-GAL4 and Ey-GAL4 lines were used to broadly express UAS-transgenes in the eye (Wernet et al., 2003). The FLP/FRT (Xu & Rubin, 1995) and MARCM techniques (Lee and Luo, 2001) were used to generate genetically marked clones by FLP-mediated mitotic recombination. We used the <u>E</u>yeless-<u>G</u>AL4/<u>U</u>AS-<u>F</u>LP (EGUF) technique, by which the retina is rescued from Hid-mediated apoptosis if mutant cells can create viable tissue, to generate entirely mutant eyes (Stowers and Schwarz, 1999). The FLP-Out/GAL4 technique was used to express desired transgenes in genetically marked clones. *y w* Ey-FLP; Ubi-mRFP, FRT40A was used to generate *pten* or *Pi3K-reg* mutant FLP/FRT clones, Ey-FLP, UAS-GFP; tub-GAL80 FRT40A; Ubi-GAL4 to generate MARCM clones and GMR-Hid FRT40A; Ey-GAL4, UAS-FLP to generate entirely mutant eyes. Mitotic and FLP-out clones were induced by a heat shock for 30 minutes at 34°C.

### Molecular biology and construction of genetically encoded reporters

We generated several new reporters driven by the ubiquitin promoter including Ubi-GFP::Pten2, Ubi-Pi3K21B::GFP, and Ubi-GFP::Abi, and several UAS-driven transgenes including UAS-GFP::Pten2, UAS-GFP::Pten (C132S), and UAS-GFP::Pten (G137E) and UAS-PI3K21B::GFP. The *pten* open reading frame (ORF) was amplified from cDNA clone IP16020 (RRID:DGRC_1603573), the Pi3K21B ORF from LD42724 (RRID:DGRC_5517), and the Abi ORF from LD37010 (RRID:DGRC_2550). The PCR products were inserted into pENTR plasmids by TOPO cloning. All expression clones were generated by the Gateway technology (Walhout et al. 2000) using the *Drosophila* Gateway Vector Collections (gift of T. Murphy and C.-Y. Pay) using the Clonase II reaction to fuse the ORFs in frame with a desired fluorescent protein. Site-directed mutagenesis using Agilent Technologies QuickChange II was performed to mutate Pten by substituting the cysteine at position 132 to serine to generate UAS-GFP::Pten2^C132S^ or by substituting the glycine at position 137 to glutamine to generate UAS-GFP::Pten2^G137E^. Ubi-Pi3K21B::GFP was inserted at the P{CaryP}attP2 site (estimated at 68A4) by PhiC31-mediated recombination. We introduced an AttB site to pUGW using site-directed mutagenesis using a New England Biolabs NEB Q5 Site-directed mutagenesis kit (Akbari et al., 2009). Transgenic fly lines carrying these transgenes were established by standard methods by BestGene, Inc.

### Method detail

#### Immunofluorescence

White prepupae (0h APF) were selected and aged on glass slides in a humidifying chamber at 25°C. Pupal eyes were dissected in phosphate-buffered saline, fixed for 35 minutes in 4% paraformaldehyde in PBS, and stained with antibodies in PBS with 3% BSA, 0.3% Triton X-100, and 0.01% Sodium Azide. Primary antibodies used were rat anti-E-cad (DSHB #DCAD2, 1:100), mouse anti-Dlg (DSHH #4F3, 1:500), and guinea pig anti-Sdk (Astigarraga et al., 2018). Alexa647 and Alexa488 (Molecular Probes) and Cy3 and Cy5 (Jackson ImmunoResearch) conjugated secondary antibodies were used at 1:150.

#### Sample preparation for live imaging

Pupae were selected and aged as described above. Larvae and pupae expressing UAS-Rac1 were aged for 48h at 18°C and then shifted to 25°C (for quantification of polarity) or 30°C (for qualitative observations of ruptures in GFP::Abi, Utr::mCh eyes) for 4 hours before imaging to conditionally increase transgene expression. Live imaging of pupal eyes was conducted as previously described (Del Signore et al., 2018). Prior to imaging, the operculum and surrounding pupal case were peeled carefully to expose the eyes. Pupae were inserted in a slit created in an agarose block with one eye facing the coverslip. The agarose block was fitted with a custom-made rubber gasket and capped with a custom-built humidified chamber.

#### Confocal imaging

Image data were collected on a Zeiss LSM800 laser scanning confocal microscope with Airscan. For analysis of epithelial remodeling and protein dynamics, an image stack was obtained every 1 minute unless otherwise noted with pinhole size=1 AU using a 63X, 1.4 NA, plan Apochromat oil immersion objective, with a laser intensity of 0.2-1%, 0.7 μm per optical section with a 10-50% overlap between sections, using a bidirectional scan at a speed of 7 without averaging.

#### Laser nano-ablation

Laser nano-ablations to measure bond tension were performed using a Zeiss LSM880 NLO with a near-infrared InSight X3 multi-photon tunable laser (680-1300nm) using a 740nm multiphoton excitation. Samples were imaged with α-Cat::Venus to identify the junctions for ablation using a 63X oil immersion objective (NA 1.4). A field size of 1x8 pixels was ablated in the center of LC-LC contacts using a 30-40% laser output, at a scan speed of one, with one iteration.

Images were collected every 0.5 seconds for 120 seconds and ablations were performed 3 seconds after the beginning of each experiment. To determine the initial recoil velocity (V_max_), we measured the distance between vertices of the ablated cell-cell contacts in Fiji and plotted it as a function of time using Prism 9 (GraphPad). V_max_ was determined using a fitting function f(t) as follows: f(t) = A · (1-e^-t/τ^) and V_max_ = df(0)/dt = A/τ where A is the total retraction amplitude and *τ* the decay rate constant.

### Quantification and Statistical Analysis

#### Quantitative assessment of eye phenotypes

We processed images and counted features using Fiji. We created Regions of Interest (ROIs) in the center of retinas in which to count cellular defects. Then we used the Cell Counter tool to count defects within the area. In addition, we counted the total number of edges, vertices, and ommatidia within the ROI. The following defects were normalized to the number of edges: 1) A missing LC, defined as any edge with only one bristle and one lattice cell, or two lattice cells and no bristles. 2) A separation, defined as any edge along which 1° cells from two different ommatidia were in direct contact. 3) An extra LC, defined as any edge with more than three lattice cells, or two bristles and more than two LCs. The following defects were normalized to the number of vertices: 1) A rosette, defined as a bristle with four LCs contacting it instead of the normal three. 2) A missing 3° LC, defined as a vertex at which a 3° LC was missing and where a contact between multiple secondary cells was formed. 3) A misplaced bristle, defined as an edge with a bristle in the middle. Statistical differences between conditions were examined using a Chi-Square Test. For counting numbers of rosettes in FLP/FRT and MARCM clones, a vertex was considered WT if the vertex and all adjacent secondary cells were WT, mutant if all cells were mutant, and mixed otherwise.

#### Analysis of contact dynamics

We examined contact dynamics at ∼28h APF at the end of the pruning stage, when some, but not all, ommatidial edges had 2 or more 2° cells. We processed images in ZEN 2.6 and Fiji, using ZEN’s Z-Stack Alignment method and applying a Gaussian blur (sigma=1.3). We took maximal intensity projections of 3-5 slices for images featuring GPH and Rac1::GFP and of all 10 slices for other reporters. Individual LC-LC contacts along horizontal edges were traced manually at 1-minute intervals for 60 minutes using a Fiji plugin with a line segment selection producing 40-60 ROIs per contact. A custom Fiji “Junction Analyzer” script measured the length of each ROI. Then, a Python script operating on the extracted length values calculated the features of the pulsing. The data were smoothed using a Gaussian filter with sigma=2. Pulse amplitudes were taken using the absolute value of the difference between adjacent local maxima and minima. This value was divided by the average contact length to get the normalized amplitude. Pulse frequencies were taken using the number of minutes between maxima. Values for a given contact were averaged to produce an average amplitude or frequency per contact. Statistics were performed on average amplitudes and frequencies per contact using a Kruskal-Wallis test with Dunn’s Multiple Comparison test in Prism (Version 9.5.0).

#### Time-shifted Pearson’s correlation analysis

Contact dynamics were analyzed by manual tracing with a line segment selection (width=5px). Vertices were automatically placed at both ends of these line segments as circular ROIs (diameter=6px). Contact length, pixel mean, and total intensity and background for each channel at each time point were collected using the Junction Analyzer Fiji macro. For Sqh::GFP, Rac1::GFP, and GFP::Pten2, and Grp1-PH::GFP (GPH), analysis was based on the mean intensity over the full contact length, while for Utr::GFP, GFP::Abi, and Pi3K-reg::GFP, it was based on the mean intensity at the vertices. Time-resolved Pearson’s cross-correlations (time windows from +/-19 minutes) were compared between the mean fluorescent marker intensities and contact length using 40-60 min long movies and calculated using a custom Python script along with packages NumPy and SciPy. Data presented are the average Pearson’s correlations of individual contacts as noted in figure legends and are presented as the mean correlation (R) +/- SD. A one-sample t-test was used to calculate whether average Pearson R-values and time shifts were significantly different from zero. To calculate whether the peak average R-value differed from the average R-value at a time shift of 0, a paired t-test was used. An unpaired t-test was used for comparisons between groups to analyze normally distributed data, while the Mann-Whitney *u* test was used to analyze non-normally distributed data.

#### Quantification of contact polarity in Rac1 overexpression eyes

We processed images in ZEN and Fiji as described above, with multiple maximal intensity projections of 3-5 slices taken for each image to focus on different patches of cells. Levels of reporters at tAJs were measured in Fiji by averaging the mean intensity at two circular ROIs of size 12, one placed at each vertex. Levels of reporters at bAJs were measured using a single circular ROI, defaulting to size 12 but reduced in size if necessary to avoid overlapping with the tAJ ROIs. We then took the ratio between the intensity at tAJs and bAJs. Comparisons between genotypes were performed with an unpaired t-test for normally distributed data and a Mann-Whitney *u* test for non-normally distributed data.

## SUPPLEMENTARY MATERIALS

**Supplemental Figure 1:**
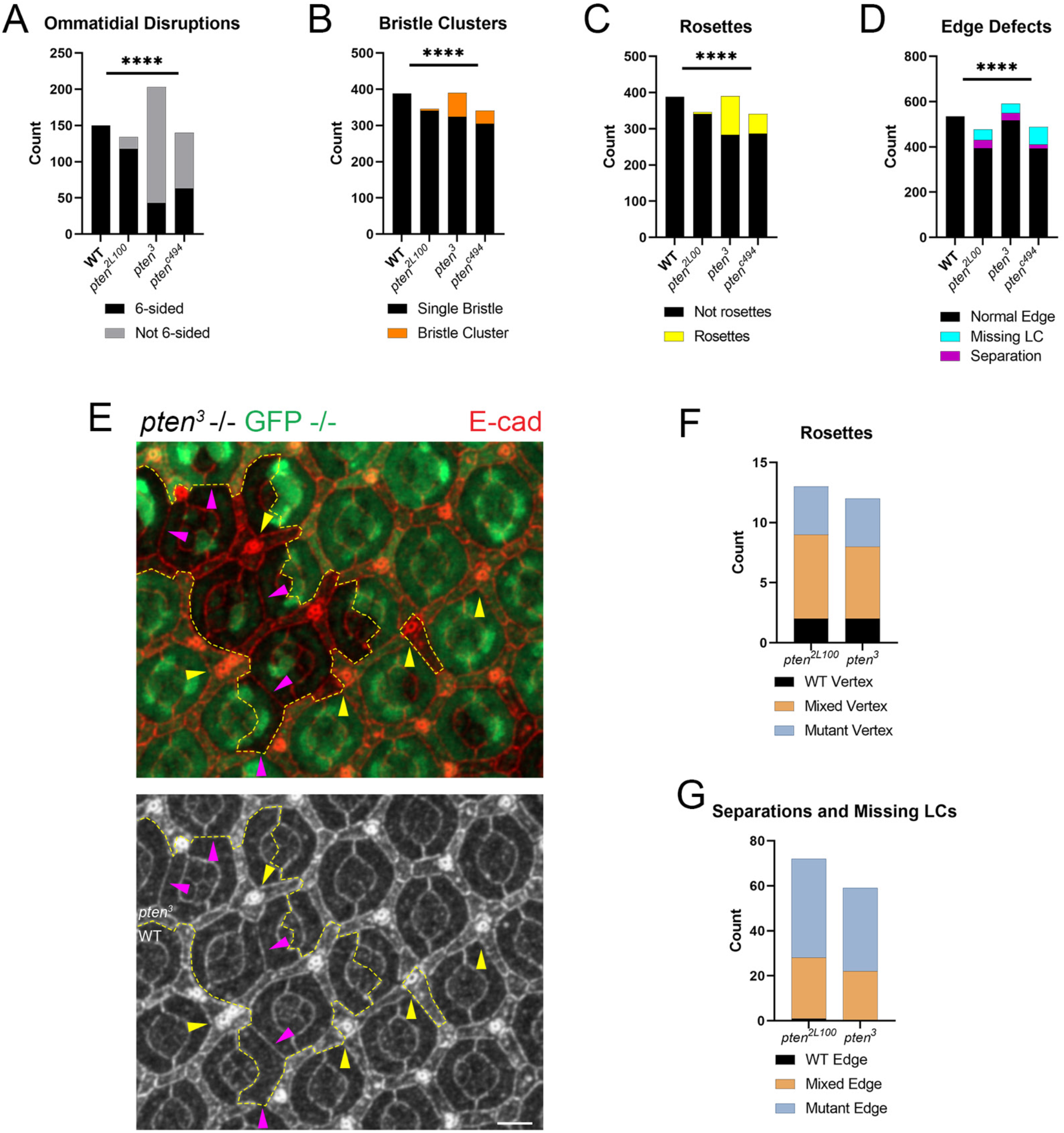
Quantification of lattice defects in *pten* mutant eyes and clones compared with WT. (A-D) Quantification of counts of specific defect types in the *pten* mutants compared with WT. *pten* mutant eyes exhibit more (A) ommatidia that are not six-sided, (B) vertices with multiple bristles, (C) cellular rosettes and (D) edges missing LCs and/or exhibiting aberrant 1°-1° contacts. χ^2^ tests; ****p < 0.0001. (E) A negatively marked *pten* mutant clone generated using the FLP/FRT system (outlined in yellow). Yellow arrowheads mark cellular rosettes; purple arrowheads mark aberrant 1°-1° contacts. (F-G) Quantifications of the mutant and WT cell composition of (F) rosettes and (G) aberrant edges in eyes containing *pten^2L100^* and *pten^3^*mutant clones compared with WT.

**Supplemental Figure 2:**
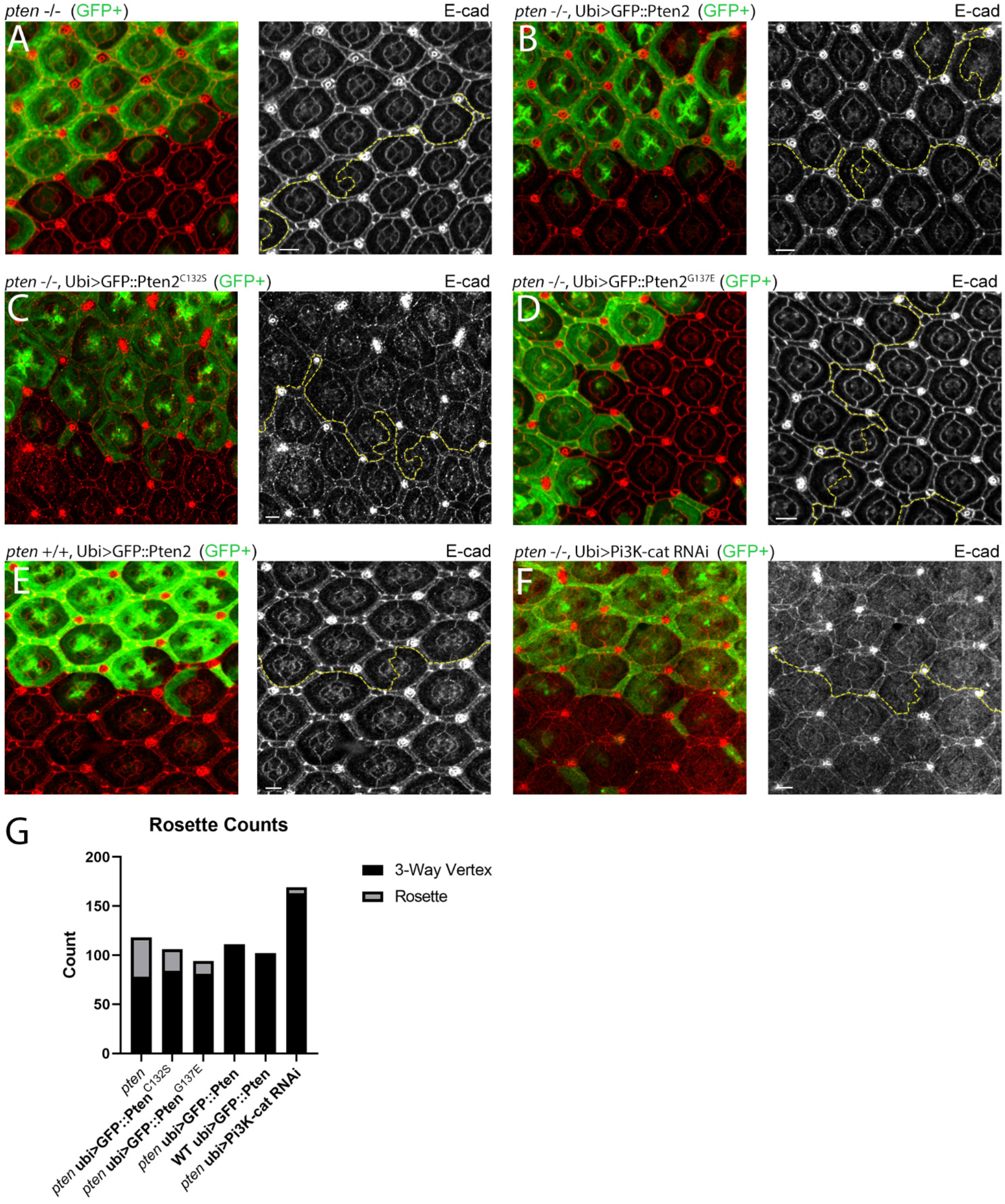
Pten lipid phosphatase activity is required for epithelial remodeling. (A-D) GFP-marked *pten* clones in a background expressing (A) no transgene, (B) WT Pten, (C) Pten^G137E^ deficient for protein and lipid phosphatase activity, and (D) Pten^C132S^ deficient for lipid phosphatase activity only. (E) A GFP-marked clone in a WT background expressing WT Pten as a positive control. (F) GFP-marked *pten* clones with RNAi-mediated depletion of Pi3K-cat. (G) Counts of cellular rosettes in clones of the above five genotypes as a proxy for overall lattice disorganization. WT Pten accomplishes a nearly complete rescue of the *pten* phenotype, while the G137E and C132S variants provide little to no rescue. RNAi depletion of Pi3K-cat accomplishes a partial rescue as well.

**Supplemental Figure 3:**
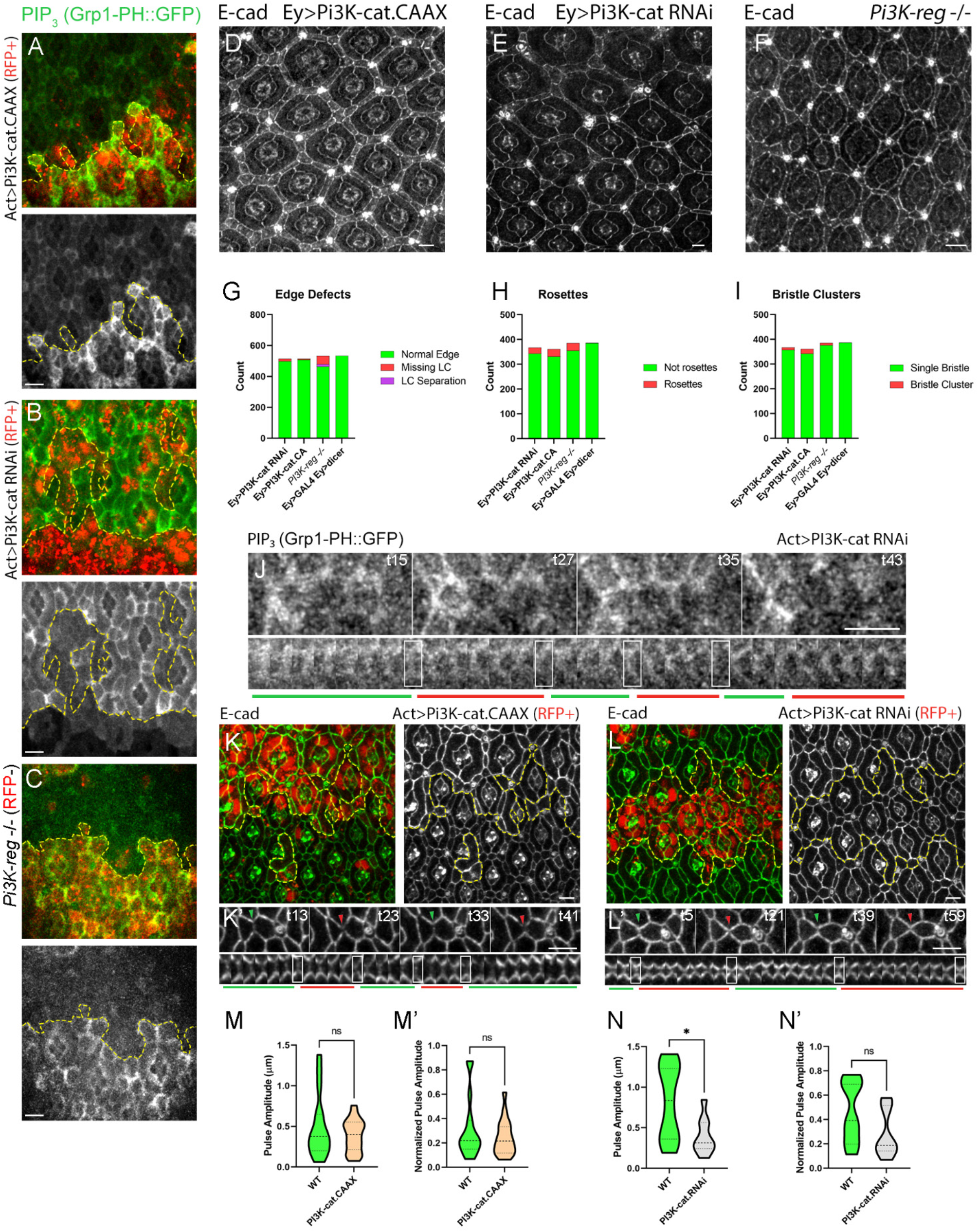
Effects of Pi3K on lattice remodeling and PIP_3_ levels. (A-C) 28h APF clones in three genotypes affecting Pi3K function, all expressing tubulin-GPH. (A) Act>Pi3K-cat.CAAX (B) Act>Pi3K-cat.RNAi (C) *Pi3K-reg* -/-. (D-F) Cellular defects in three genotypes affecting Pi3K function, 34-38h APF. (A) Ey>Pi3K-cat.CAAX. (B) Ey>Pi3K-cat.RNAi. (C) *Pi3K-reg* mutant eye. (G-I) Quantification of counts of specific defect types in the above genotypes compared with WT. Both overexpression and depletion of Pi3K function increase the numbers of (G) edges missing LCs, (H) vertices with multiple bristles, and (I) cellular rosettes. (J) Dynamics of PIP_3_ in a clone expressing Act>Pi3K-cat RNAi. In addition to overall low levels, PIP_3_ pulsing is reduced. (K-L) Clones expressing E-cad::GFP and (K, K’) Act>Pi3K-cat.CAAX or (L, L’) Act>Pi3K-cat.RNAi, displaying (K, L) cellular defects and (K’, L’) reduced LC-LC contact length pulsing. Quantification of (M-N) absolute and (M’-N’) normalized pulse amplitudes in clones expressing (M) Pi3K-cat.CAAX and (N) Pi3K-cat RNAi compared with adjacent WT regions. Unpaired t-test, n=9-18 contacts *p<.05.

**Supplemental Figure 4:**
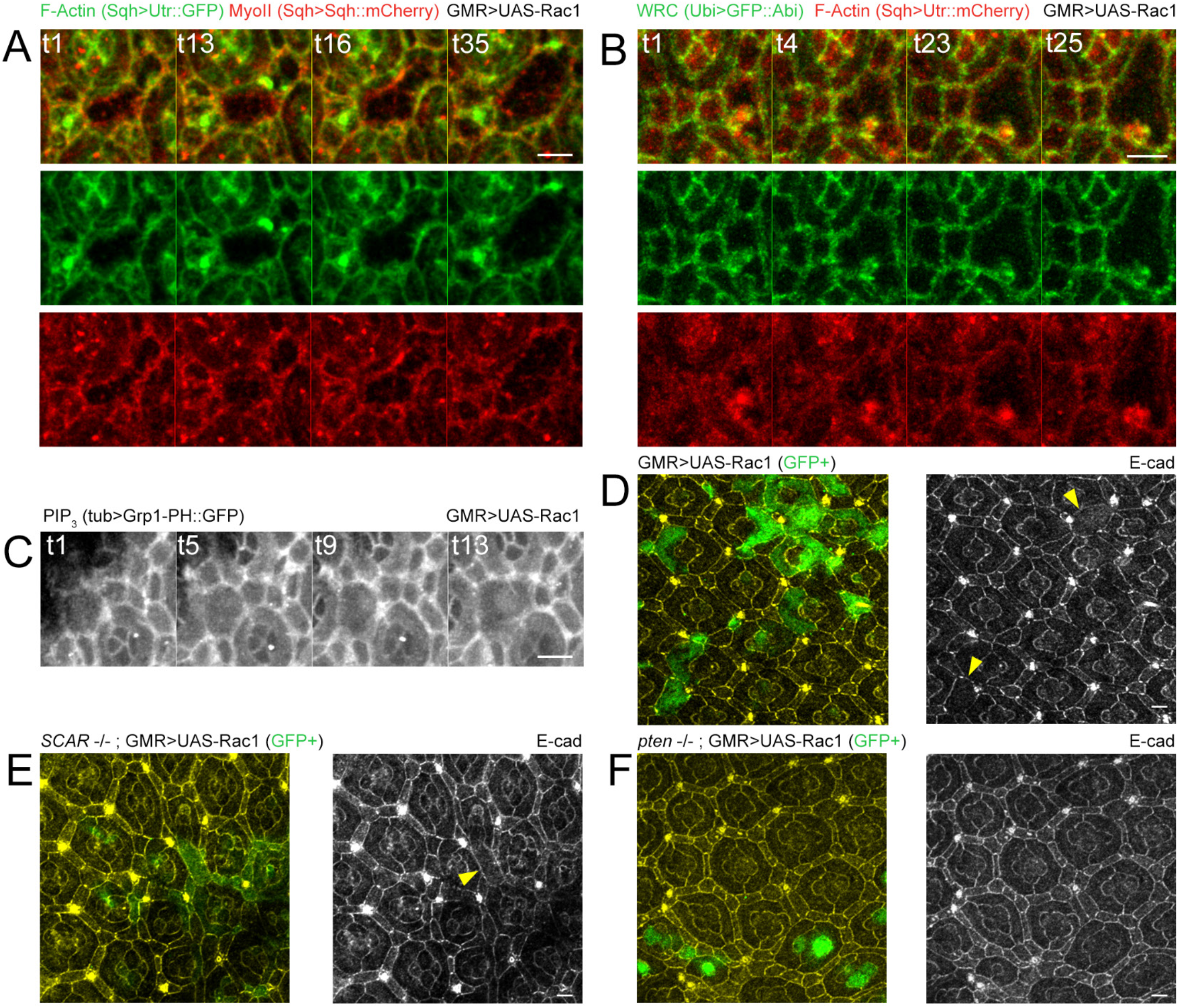
Large-scale tissue disruptions in Rac1-overexpressing eyes. (A) Snapshots of an expanding tissue rupture. Expansions of the rupture are accompanied by F-actin bursts, while MyoII extends into the hole formed by the rupture. (B) Snapshots of another rupture showing enrichment of both Abi and F-actin at its edges as they expand. (C) Snapshots of a rupture with labeled PIP_3_ enriched at breaking contacts. (D) GFP-marked clones overexpressing Rac1. Arrowheads mark ruptured cells. (E) GFP-marked clones overexpressing Rac1 in a *SCAR* -/- background. An arrowhead marks a ruptured cell. (F) GFP-marked clones overexpressing Rac1 in a *pten* -/- background. There are no ruptured cells and few surviving mutant LCs, though non-autonomous defects persist in surrounding ommatidia.

**Supplemental movie 1: Cellular dynamics in WT compared with *pten* mutant eyes.** (Corresponds to Fig. 2)

Left: Time-lapse of a WT eye showing normal dynamics of repeated cell contact expansion and contraction during lattice remodeling (white arrowhead). When a LC is pruned from the lattice, contact between adjacent cells is reestablished immediately (yellow arrowhead). Right: A *pten* mutant eye with a disordered ommatidium demonstrating pinched contacts that fail to expand and contract (white arrowhead) and separations between LCs following cell pruning that take up to an hour to repair (yellow arrowhead).

**Supplemental movie 2: F-actin and MyoII dynamics in WT compared with *pten* mutant eyes.** (Corresponds to Fig. 3) Far left: Time-lapse of a WT eye showing dynamics of F-actin. Note F-actin accumulation while the contact is expanding and disassembly during contraction. Middle left: Time-lapse of a *pten* eye showing F-actin dynamics.

Middle right: Time-lapse of a WT eye showing dynamics of MyoII. Note accumulation of MyoII in contracting contact and lower MyoII levels in expanding contact. Far right: Time-lapse of a *pten* eye showing dynamics of MyoII. MyoII appears in more clearly defined cables than in WT but does not accumulate and dissipate along with contact expansion and contraction, respectively.

**Supplemental movie 3: WRC dynamics in WT compared with *pten* mutant eyes** (Corresponds to Fig. 4)

Left: Time-lapse of a WT eye showing dynamics of the WRC. Note accumulation while the contact is expanding and dispersal during contraction. Right: Time-lapse of a *pten* eye showing dynamics of the WRC.

**Supplemental movie 4: PIP_3_ dynamics in WT compared with *pten* mutant eyes** (Corresponds to Fig. 5)

Left: Time lapse of a WT eye showing dynamics of PIP_3_ with E-cad for cell outlines. Note accumulation while the contact is expanding and dispersal during contraction. Center: The same ommatidium with only PIP_3_. Right: Time-lapse of a *pten* eye showing PIP_3_ dynamics.

**Supplemental movie 5: PIP_3_ dynamics in clones with increased or reduced Pi3K-cat activity.** (Corresponds to Fig. 6) Left: Time lapse of PIP_3_ dynamics in a clone expressing Pi3K-cat.CAAX (marked in red) and adjacent WT region. Right: Time lapse of PIP_3_ dynamics in a clone expressing Pi3K-cat RNAi (marked in red) and adjacent WT region. Details show close-ups of rows of clone cells.

**Supplemental movie 6: Dynamics of cells and regulators of protrusion in *Pi3K-reg* mutant eyes** (Corresponds to Fig. 6) Left: An ommatidium from a *Pi3K-reg* mutant eye in which LC-LC contacts fail to expand and contract. Center: Time lapse of F-actin dynamics in a *Pi3K-reg* mutant eye. Right: Time lapse of WRC dynamics in a *Pi3K-reg* mutant eye.

**Supplemental movie 7: Rac dynamics in WT compared with *pten* mutant eyes** (Corresponds to Fig. 7)

Left: Time lapse of a WT eye showing dynamics of Rac1. Note accumulation while the contact is expanding and dispersal during contraction. Right: Time-lapse of a *pten* eye showing dynamics of Rac1.

**Supplemental movie 8: Abnormal dynamics of F-actin, the WRC and PIP_3_ in eyes overexpressing Rac1** (Corresponds to Fig. 7)

Dynamics of F-actin (Left), the WRC (Center) and PIP_3_ (Right) in eyes overexpressing Rac1. Fluctuations of contact length are reduced, and all three molecules remain overly concentrated at contact vertices.

**Supplemental movie 9: Tissue ruptures in eyes overexpressing Rac1** (Corresponds to Fig. 7)

Top: Tissue rupture expanding in an eye with labeled F-actin and MyoII. Middle: Tissue rupture forming and expanding in an eye with labeled Abi and F-actin. Bottom: Tissue rupture forming in an eye expressing tagged Grp1-PH to label PIP_3_.

**Supplemental movie 10: Dynamics of Pi3K-reg and Pten** (Corresponds to Fig. 8)

Left: An ommatidium in an eye expressing tagged Lifeact to detect F-actin and Pi3K-reg. Note accumulation of Pi3K at vertices in expanded contacts and in dimmer spots adjacent to vertices in contracted contacts. Right: An ommatidium in an eye expressing tagged Lifeact and Pten.

